# Antibiotics Change the Growth Rate Heterogeneity and Morphology of Bacteria

**DOI:** 10.1101/2024.08.27.609914

**Authors:** Morten Kals, Emma Kals, Jurij Kotar, Allen Donald, Leonardo Mancini, Pietro Cicuta

## Abstract

A better understanding of the system-level effects of antibiotics is necessary to fight the rise of antibiotic resistance. Utilising the Multipad Agarose Plate (MAP), we monitor the growth rate and cell morphology of three clinically relevant species (*E*.*coli, S*.*aureus* and *P*.*aeruginosa*) after exposure to 11 different concentrations of 13 antibiotics (for a total of 24 microbe-antibiotic combinations). As the drug dose approaches the MIC and regardless of the mode of action, our results show a consistent increase in growth rate heterogeneity. Remarkably, drugs that affect protein synthesis consistently show the opposite trend, reducing heterogeneity. We hypothesize that growth rate heterogeneity under antibiotic treatment might therefore depend on the functional distance of the target from ribosomal activity, which is key in determining growth rate. Low heterogeneity is desirable from a clinical perspective, as the opposite is often associated to persistence and antibiotic survival. For all of the antibiotics and species tested, we also find a striking and non-trivial correlation between morphological alterations and growth inhibition. This observation allows us to introduce a new morphological parameter, MOR50, that enables the estimation of minimum inhibitory concentration (MIC) for antibiotic susceptibility testing (AST) with a single snapshot after 2.5 hours of incubation. In addition to introducing a novel, resource-efficient, rapid AST method, our findings shed new light on the effects of antibiotic perturbations on bacteria at the system level that might inform treatment design.

## 1 Introduction

Antibiotics interact with specific molecular targets within bacterial cells to inhibit essential processes. These disruptions initiate cascades of events that eventually lead to two phenotypes: growth arrest and death [1]. Although the specific molecular targets of most antibiotics are well characterised [2], the indirect processes that ultimately impair cellular growth are complex and not fully understood. A comprehensive understanding of the system effects of antibiotics would shed light on the inner workings of bacteria and could improve the rational design of treatment. Cell size is one of the properties that bacteria regulate at the system level. It is the complex, intensively studied, and still not fully understood result of the coordination of the rates of synthesis of the cell’s envelope, its DNA and its many other cytoplasmic contents. These rates in turn depend on ribosome activity and metabolism [3]. For *E*.*coli* in balanced growth, cell size has a strong positive correlation with growth rate and ribosomal content [3, 4]. Perturbations, such as those caused by antibiotics, can decouple such relations. For example, decreased DNA replication rates can lead to increased cell sizes [3], while partial inhibition of the ribosomal pool enlarges cells in nutrient-poor medium and shrinks them in rich medium [4]. However, the field does not have a holistic picture of how antibiotics impact cell morphology, especially where such changes would be linked to broader system-level effects, rather than as a direct consequence of the antibiotic target being inactivated.

In bacteria, growth rate is also controlled at the system level. When measuring antibiotic efficacy, an altered growth rate is typically the easiest phenotype to observe: the minimum concentration of antibiotic that leads to growth inhibition, known as the minimum inhibitory concentration (MIC), has guided clinical research and diagnostics for decades [5]. Several other aspects of growth can be studied to gain insight into the action of antibiotics. Sub-inhibitory growth rate changes due to antibiotics, for example, have helped shed light on important questions concerning how cells grow and regulate their macromolecular composition [6], and how they respond to perturbations [7]. The relationship goes both ways, and the cell’s growth rate can in turn alter antibiotic efficacy [8, 9]. Important bactericidal antibiotic classes, such as aminoglycosides, fluoroquinolones and beta-lactams, are essentially inefficacious against non-growing cells [10, 11]. Complex systems such as bacteria are intrinsically heterogenous [12] and bacterial populations simultaneously harbour fast, slow, and non-growing cells [13]. Heterogeneity in a genetically identical cell population is an important evolutionary trait, and it is the basis of the observed persistence to many antibiotic treatments [14]. Persister cells with reduced growth rates have been extensively studied in vitro [15] and are thought to drive infection relapses *in vivo* [16]. Interestingly, some types of persistence are thought to be triggered by exposure to antibiotics, an unwanted effect from the point of view of therapy [14].

Some of these system-level effects have now begun to be revealed. For example, our lab has demonstrated that an antibiotic such as rifampicin, which targets RNA polymerase [17], can reduce macromolecular crowding [18, 19]. This reduction is a severe, system-level perturbation that impacts the rates of nearly all cellular biochemical processes. We observed similar effects with ciprofloxacin, a fluoroquinolone that interacts with DNA gyrases [19]. Beta-lactam antibiotics, that compromise cell wall integrity, are also known to cause toxic system effects by promoting a futile cycle of cell wall precursors that depletes cellular resources [20]. Aminoglycosides, which interact with the ribosome to cause translation errors [21], are also thought to disrupt cell integrity [22], impacting the cell’s energy production mechanisms [23]. Universal system-level bactericidal strategies based on the production of reactive oxygen species (ROS) have been proposed [24], spurring debate [25, 26, 27, 28], and finding a certain degree of experimental support [29, 30, 31]. Whether by means that are ultimately universal or not, it is clear that antibiotics interact with the complex system that is the cell, introducing perturbations that can be observed in molecular pathways far beyond where they started.

We recently presented a new experimental platform that enables the high-throughput optical imaging of live microbes across different environmental conditions, the Multipad Agarose Plate (MAP) [32]. In this work, we leverage and extend the MAP to perform a systematic investigation of the growth dynamics and morphological responses of three clinically relevant bacterial species (*E*.*coli, P*.*aeruginosa* and *S*.*aureus*). We probe these bacteria out of balanced growth, subjecting them to a panel of 13 antibiotics at a range of concentrations and capturing sub-MIC, MIC and post-MIC behaviour. The platform and the accompanying analysis pipeline allow the extraction of single cell and colony parameters directly from images taken with brightfield illumination, so that samples can be completely label-free. Our results reveal that, approaching the MIC, antibiotics systematically increase the heterogeneity of growth rate across microcolonies, potentially suggesting that a persister-triggering effect might be intrinsic to several antibiotic classes. Notably, this appears to be true independently of the bacteriostatic or bactericidal nature of the antibiotic, for example the bacteriostatic rifampicin has the same effect as the bactericidal beta-lactams on *E*.*coli*. Antibiotics that target protein synthesis are a consistent exception across the different species, having the opposite effect. Again, this seems to be independent of growth stasis or death, as for example the bactericidal kanamycin shows the same behaviour as the bacteriostatic chloramphenicol.

Growth rate heterogeneity is not the only factor that shows a striking correlation with the MIC. Our investigation of single-cell morphologies across the 13 antibiotics and three microbial species not only shows that all antibiotics cause morphological alterations, but unveils a strong correlation between the degree of morphological alteration and the MIC. Using dyes, several works have previously linked antibiotic efficacy [33] and even mechanism of action [34] with morphology using single-cell imaging. Using our insight on label-free morphology alone, we introduce a simple metric that we call *morphological change 50* (MOR50). MOR50 is the antibiotic concentration that, after 2.5 hours of incubation with exponentially growing cells, induces a morphological change of 50% of the largest morphological change induced at any concentration of that antibiotic in said cells. We show that MOR50 can be used to reliably determine the MIC from a single snapshot (fig. 1.).

**Figure 1.**
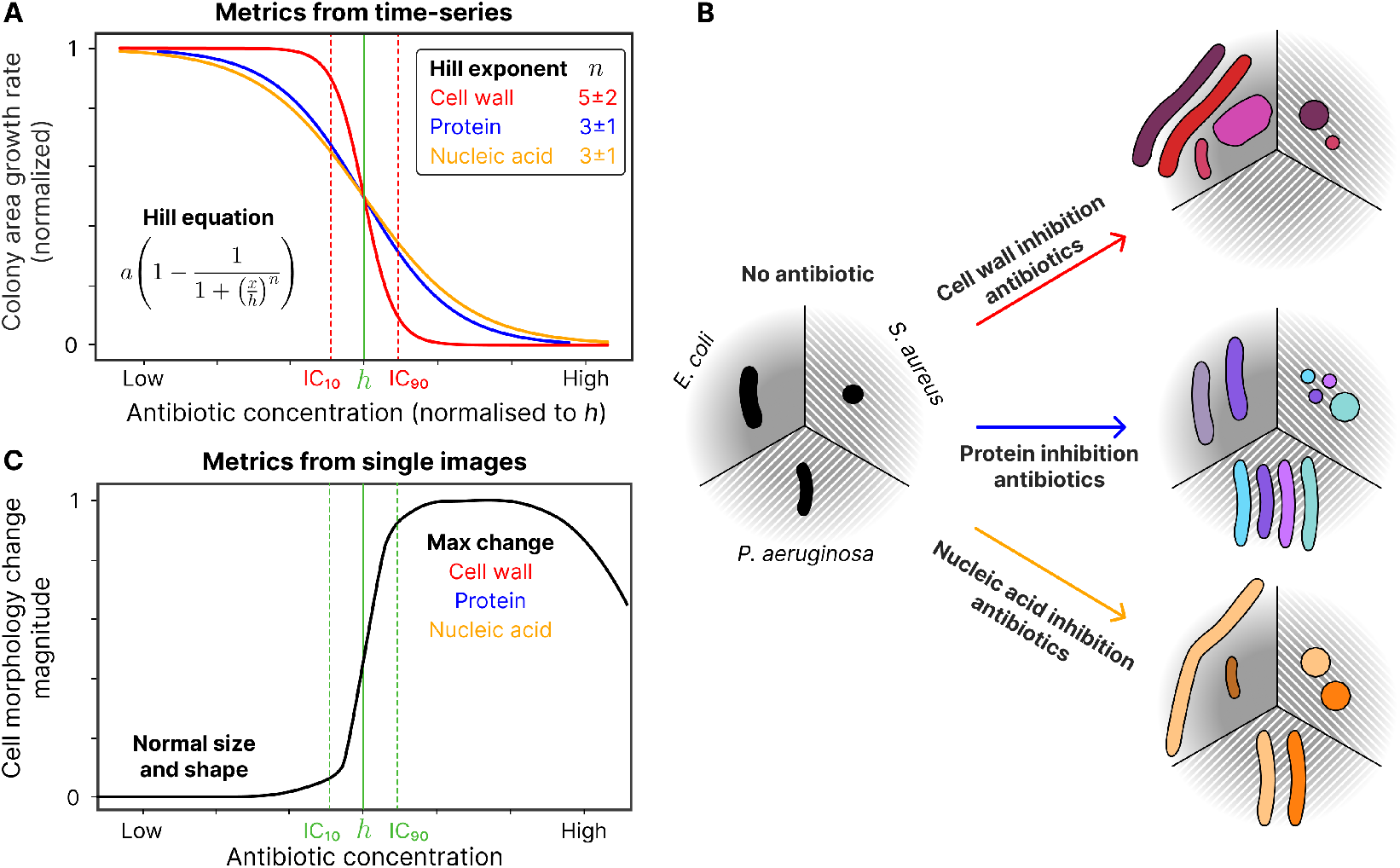
Our approach combines and shows correlations between observations over time (growth rate, A) and single-time assessment of the morphology of individual cells, C. **A** Hill fits are performed independently for each antibiotic, species and repeat to determine the relationship between antibiotic concentration and bacteria growth rate. The Hill curve is characterized by three parameters: the maximum growth rate *a*, the Hill fit coefficient *h*, and the Hill fit exponent *n*. We report MIC as the concentration where the growth rate is reduced by 90%, inhibitory concentration (IC) *IC*_90_. The Hill fit coefficient *h* is the concentration of antibiotic that reduces the growth rate by 50% *IC*_50_, and the *IC*_10_ line is also indicated for reference, with examples of *IC*_90_ and *IC*_10_ only shown for the cell wall synthesis inhibitors. The Hill fit exponent *n* is a measure of how steep the drop is. The cell wall synthesis inhibitors generally produce fits with the largest *n* values. The protein and nucleic acid synthesis inhibitors have significantly smaller *n* values, indicating a larger concentration range that inhibits growth without completely stopping it. The value of *h* is well known to link to the MIC as measured in other assays. Lines are plots of the Hill equation with the values given in the panel, which are discussed in the text. **B** Schematic showing how the different antibiotics impact cell morphology for the three tested species of bacteria. The black cell masks represent the typical morphologies of the bacteria species without an antibiotic present, whereas the coloured bacteria to the right represent the morphological changes induced by that antibiotic (colour-coded consistently with fig. 2). The cell wall synthesis inhibitors generally produce the largest change in the cell area, whereas the protein synthesis inhibitors produce the smallest changes. Some antibiotics induce an increase in size; others cause a decrease. Data for this plot is shown in fig. 5. **C** Schematic plot showing that morphological changes depend on concentration. As shown in **B**, there are large differences in the amplitude of changes, but normalization is possible and makes all antibiotics produce a similar general pattern, as shown here, regardless of the mechanism of action. The largest changes in morphology occur when the growth rate is significantly reduced, at antibiotic concentrations that we show are related to the MIC.

Taken together, our results identify an extensive and un-appreciated tendency towards heterogeneous growth among pathogens when exposed to sub-inhibitory concentrations of antibiotics that do not directly inhibit protein synthesis. We expect that this will motivate further research aimed at understanding whether such phenomena may lead to the formation of persisters, and whether their formation relies on mechanisms that require protein synthesis.

We further argue that our new metric, MOR50, could pave the way for a new generation of rapid phenotypic AST methods, potentially transforming the landscape of microbial diagnostics and antibiotic therapy.

## 2 Results

### 2.1 The MAP platform accurately measures the MIC of *E*.*coli, P*.*aeruginosa* and *S*.*aureus* based on growth rate

Leveraging the recently developed MAP platform [32], we first set out to assay the MIC of a panel of 13 antibiotics against *E*.*coli, P*.*aeruginosa* and *S*.*aureus*, three human commensals that can turn pathogenic. This was done based on the tracking of growth rate over the first three hours after exposure to antibiotics fig. S4-S6. As previously [32], we used Hill curves to model growth rate as a function of antibiotic concentration fig. 2.A. See fig. S7-S9 for fits performed on each repeat for all combinations of species and antibiotics. The fit is highly time-dependent for some antibiotics because death or growth halt occurs, respectively, only after a significant amount of damage has accumulated in the cell or after the action of a significant amount of molecular targets of the antibiotic has been blocked. It is, therefore, crucial to pick the time window used for analysis carefully. We chose to use the 2.5-hour time-point for the MIC determination based on the growth rate, as by then, all of the different antibiotics had performed their action (bacteriostatic or bactericidal). The Hill exponent *n* measures the steepness of the drop between growing and non-growing conditions across the various antibiotic concentrations, with larger values indicating a steeper drop. The correspondence with EUCAST values confirmed the validity of the platform for the assay of antimicrobial susceptibility beyond *E*.*coli*, see fig. S27.

**Figure 2.**
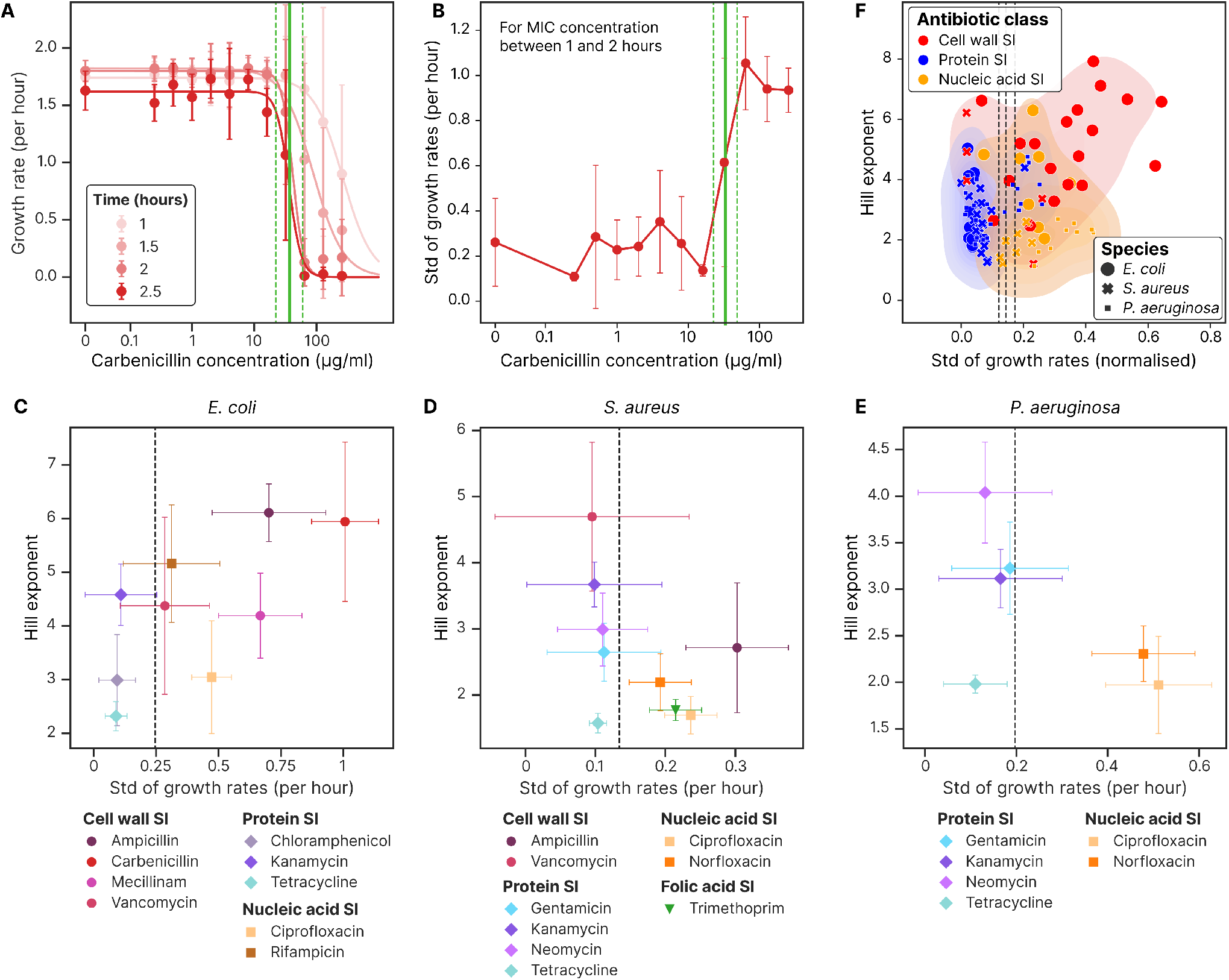
Exploring the relationship between antibiotic concentration dependence (represented by the Hill exponent *n*) and growth rate heterogeneity (GRH, here represented by std in growth rates) for different classes of antibiotics in *E*.*coli, P*.*aeruginosa* and *S*.*aureus*. **A** showing how the *E*.*coli* growth rate Hill fit for carbenicillin changes over time. Initially, the area growth rate is larger for all concentrations as the cells exposed to high concentrations of antibiotics swell up. After 2.5 hours, we see a sharp drop from viable pads to those where the antibiotic induces lysis. This 2.5-hour time-point is used for subsequent values of *n*, with corresponding values for *IC*_10_, *IC*_50_ = *h* and *IC*_90_ represented by vertical green lines. **B** The variation in growth rate on a single pad is computed and averaged across repeats in the 1 to 2.5-hour time period. We call this metric GRH. We see that the high concentrations of antibiotics produce a much higher variation in growth rate between the colonies and that this change is linked to the MIC. **C** Combining the Hill exponent *n* from **A** and GRH from **B** shows some trends in behaviour between the antibiotic classes here in *E*.*coli*. The data points represent the mean and std across repeats, with at least four repeats per condition. Each hue represents a separate antibiotic, and each symbol represents the action mechanism of that antibiotic. The vertical line signifies the mean value for the negative control, giving a benchmark for how the bacteria behave without antibiotics present. **D** Same scatterplot for *S*.*aureus*. **E** Same scatterplot for *P*.*aeruginosa*. **F** Normalizing the growth rate heterogeneity by the antibiotic-free growth rate and plotting data for all species, we see some clustering. The shaded area represents the kernel density estimate (KDE) plot of the classes, showing the smoothed 2D probability density functions. Hues represent the different classes of antibiotics, and the symbols and sizes represent the bacteria species. The vertical lines represent the normalized mean GRH for the negative controls for *E*.*coli, P*.*aeruginosa*, and *S*.*aureus*, respectively (from left to right).

### 2.2 All antibiotics except protein synthesis inhibitors increase growth rate heterogeneity approaching the MIC

Among the antibiotics considered, we noticed an interesting pattern in a metric that we call growth rate heterogeneity (GRH). The GRH is the variation in growth rate between colonies in the same growth conditions. It is computed as the standard deviation in growth rate for each pad and then averaged across repeats in the 1 to 2.5-hour time period. For all antibiotics and across the three species (fig. 2.B and fig. S10-S12), we see a consistent increase in the variation of the growth rate between colonies when the antibiotic concentration approaches the MIC. In our screening, which included a total of 24 antibiotic-species combinations with 4*±*1 repeats per antibiotic and 30*±*10 colonies per repeat, the correlation between the increase in heterogeneity and MIC was so striking that in many cases, heterogeneity alone could be used as a proxy for the MIC (fig. S13-S16). Interestingly, one class of antibiotics showed the completely opposite trend. Protein synthesis inhibitors such as chloramphenicol and tetracycline drastically reduced colony-to-colony growth rate variability approaching the MIC, again in a highly consistent way (fig. S13-S15).

We then checked whether GRH was associated with the fitting parameter *n*, as smaller changes in the delivered or perceived antibiotic concentration would give rise to larger growth rate differences. Interestingly, we found the two to be correlated in *E*.*coli*, but not in *S*.*aureus* and *P*.*aeruginosa* (fig. 2.CDE). The reason for this is that the correlation in *E*.*coli* is mostly driven by cell wall targeting antibiotics to which *P*.*aeruginosa* [35], and to a smaller extent *S*.*aureus* [36], are inherently resistant. Indeed, when we normalize the GRH by the antibiotic-free growth rate and plot data for the three species together, we see clustering between the three general mechanisms of action fig. 2.F. Protein synthesis inhibitors consistently cluster closest to the origin, cell wall synthesis inhibitors have high *n* and GRH, while nucleic acid synthesis inhibitors take an intermediate position.

### 2.3 Significant changes in morphology occur after 2.5 hours of incubation with antibiotics (six *E*.*coli* doublings and four *S*.*aureus* or *P*.*aeruginosa* doublings)

Next, we turned our attention to single-cell morphology. Using the same datasets as for the growth rate analysis, we performed single-cell segmentation on the micrographs to compute cell areas, lengths and widths. To do so, we expanded our analysis pipeline to include a classical cell segmentation step, informed by the colony segmentation. Figure fig. 3.A shows the *E*.*coli* cell segmentation masks atop brightfield microscopy data, highlighting that our analysis pipeline can appropriately segment *E*.*coli* cells that grow filamentous in response to ciprofloxacin. For comprehensive benchmarking of this algorithm on the three species, see tab. S2. In fig. 3.B, we show how this phenomenon is time-dependent. All cells start out with the same morphology when seeded on the pads, and only after some time do changes caused by the antibiotic begin to appear. It is, therefore, important to be consistent with the experimental protocol and time-point used for analysis. We show that morphological changes are concentration-dependent by taking data from the 2.5-hour period again and plotting morphology parameters against antibiotic concentration. Cells treated with ciprofloxacin only elongate in a subset of the concentrations tested, presumably because low doses are insufficient to cause changes and high doses kill the cells quickly, not giving them time to grow at all fig. 3.C.

**Figure 3.**
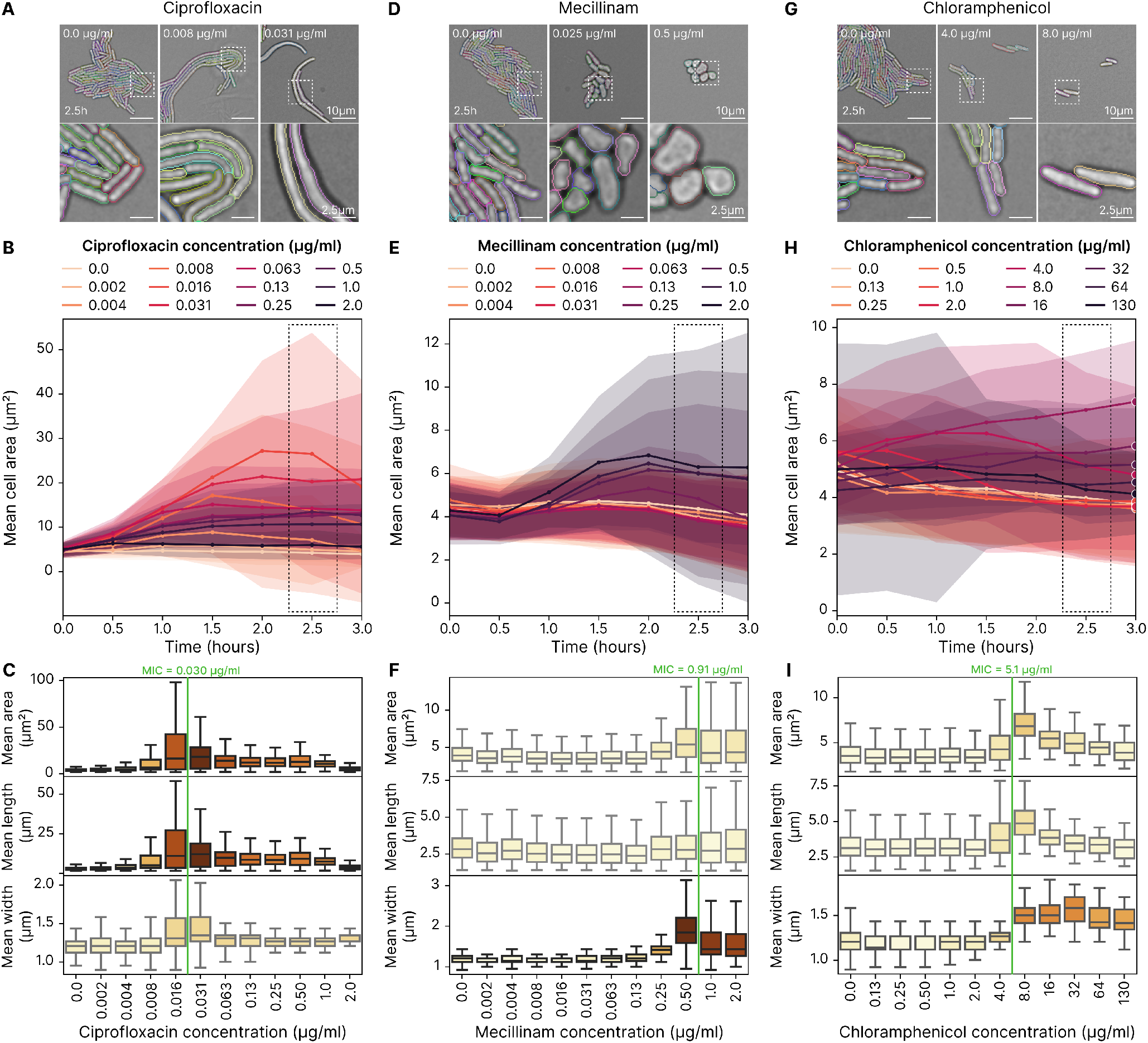
The morphology of *E*.*coli* changes significantly when exposed to antibiotics. **A** Sample frames from pads with different antibiotic concentrations highlight how the antibiotics impact cell morphology. Single-cell segmentation masks are annotated, with different colours for each cell. In this case, 2.5 hours after incubation, it is clear how the nucleic acid synthesis inhibitor ciprofloxacin induces filamentous growth. These images also show the single-cell segmentation masks, with a different colour for each cell. The bottom row is a 4 zoom of the corresponding frame in the top row. **B** This filamentous growth pattern starts to become apparent after some time, and the largest differences are seen around 2 hours of growth. Each line represents the mean area per cell for a given antibiotic concentration, and the shaded area represents the standard deviation with data originating from four repeats. Darker colours correspond to higher antibiotic concentration, and data points are binned to the nearest 30 minutes. **C** Using morphology data from 2.5 hours, it is clear how concentrations of antibiotics close to the MIC produce the largest changes in morphology. The area increases significantly, primarily linked to an increase in cell length. The MIC is indicated by the green vertical line and averaged based on the growth rate for each of the four repeats. **D** The cell wall synthesis inhibitor mecillinam causes the cells to grow into a more spherical shape. **E** It takes some time for this change to become apparent, and the largest changes are seen around 2 hours of growth. Data from four repeats. **F** Again, using morphology data from 2.5 hours, we see that the increase in width is the most pronounced change, whereas the length has hardly changed. This increase is most pronounced at concentrations close to the MIC. **G** The protein synthesis inhibitor chloramphenicol causes the cells to grow larger. **H** This change starts to occur quickly and is apparent after only an hour of growth. Data from four repeats. **I** Chloramphenicol causes both a moderate length and width increase. The area increase starts to become apparent at 4 µg mL^*−*1^, just below the MIC. The largest change is for 8 µg mL^*−*1^, the concentration just above the MIC.

The cell wall synthesis inhibitor mecillinam also causes morphological alterations in *E*.*coli*. While these manifested mainly in length for ciprofloxacin, mecillinam causes the cells to grow into spheroids fig. 3.D. These are also well-segmented by our pipeline. Like for ciprofloxacin, the change in width is time and concentration-dependent, with the largest shapes occurring after 2 hours of growth fig. 3.E and fig. 3.F.

The protein synthesis inhibitor chloramphenicol also produces an increase in cell size fig. 3.G. The increase becomes apparent only after some delay as a consequence of growth and division fig. 3.H. Like in the other cases, the change in size is concentration-dependent and caused by both increased width and length fig. 3.I.

A final example we will mention is vancomycin, a cell wall synthesis inhibitor that is the only antibiotic tested that causes *E*.*coli* cells to shrink in size. Again, the shrinkage becomes apparent only after some delay as a consequence of growth and division fig. S17. Like in the other cases, the change in size is concentration-dependent, and it is due to a reduction in length that dominates the small increase in width fig. S20.

See fig. S17-S19 for plots highlighting morphology over time for all the tested antibiotics with *E*.*coli, P*.*aeruginosa* and *S*.*aureus*. Taken together, these results suggested that, for cells growing exponentially at the time of seeding, the largest morphological changes generally occur after about 2.5 hours of growth on the MAP.

### 2.4 All antibiotics induce changes in cell morphology

Having established the 2.5-hour time point as the time of choice for assessing morphology, we investigated all of the remaining antibiotic-microbe combinations fig. 4. Images of sample microcolonies of *E*.*coli, S*.*aureus*, and *P*.*aeruginosa* (fig. 4A, C and E respectively) treated with different antibiotics are given. Remarkably, all of the antibiotics tested, albeit with different magnitudes, caused noticeable morphological changes on all of the species tested fig. 4B, D and F. With the exception of vancomycin, *E*.*coli* increased in size when treated with antibiotics, reaching the biggest volumes when treated with beta-lactams or ciprofloxacin fig. 4B. The morphological changes of *S*.*aureus* showed an almost even split between volume increase and decrease. Except for tetracycline, protein synthesis inhibitors and vancomycin tended to produce smaller cells, while DNA synthesis inhibitors and ampicillin caused cell enlargement fig. 4D. None of the antibiotics tested caused shrinkage of *P*.*aeruginosa*. Instead, this species increased in size in response to all treatments fig. 4F.

**Figure 4.**
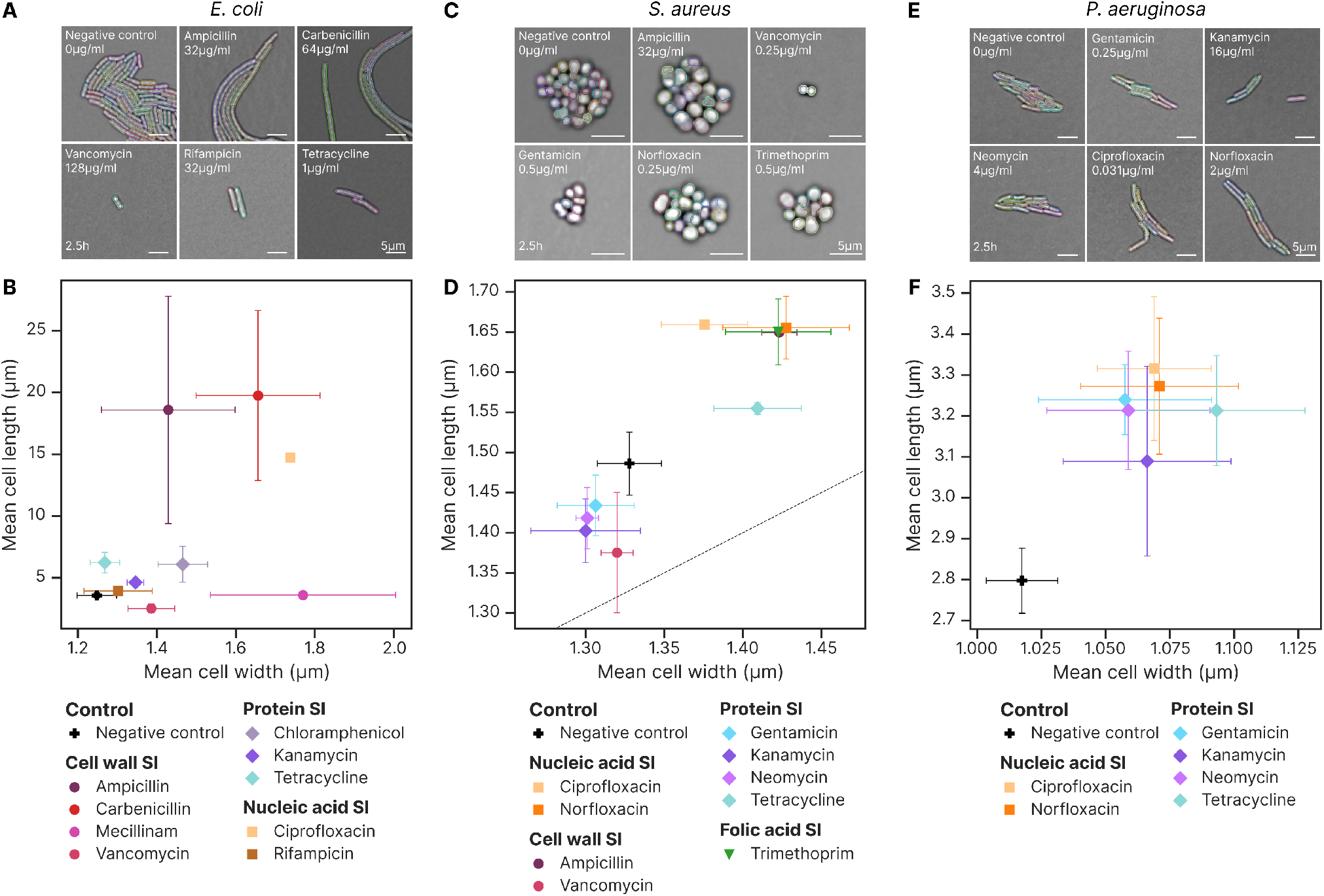
All antibiotics induce a change in morphology, though to different extents. These plots show data from the concentration of antibiotic closest to cell MIC after 2.5 hours of imaging. **A** These sample images show how *E*.*coli* responds to antibiotics from the different action mechanism classes. It is apparent that at this concentration, the biomass generated has been reduced for all antibiotics. All antibiotics produce an increase in area, although this increase is very small for rifampicin especially. See fig. 3 for images from ciprofloxacin, mecillinam and chloramphenicol. **B** Cell wall synthesis inhibitors produce large changes in morphology, as expected. However, nucleic acid synthesis inhibitors and protein synthesis inhibitors also generally produce significant changes in morphology, which is more surprising. Rifampicin is the antibiotic with the least effect on morphology. **C** Shows how *S*.*aureus* responds to a sample of antibiotics. Ampicillin, gentamicin, norfloxacin and trimethoprim produce an increase in area compared to the control, whereas vancomycin is an example of an antibiotic that induces a reduction in size. **D** Both cell wall and nucleic acid synthesis inhibitors impact *S*.*aureus* morphology significantly for all tested antibiotics. The protein synthesis inhibitors produce smaller changes. Kanamycin, neomycin, and gentamicin are approaching the noise floor of these measurements. The dashed line shows *x* = *y*, meaning perfect spheres would fall on this line. **E** Shows how *P*.*aeruginosa* responds to a sample of antibiotics. All the antibiotics induce a subtle increase in area. **F** All these antibiotics produce small but significant increases in cell length and width for *P*.*aeruginosa*.

Overall, as seen in fig. 5, cell wall synthesis inhibitors generally produced the largest changes in single cell volume, whereas the protein synthesis inhibitors produced the smallest. Some antibiotics caused consistent changes across the different species (like ampicillin, norfloxacin and ciprofloxacin), while others showed significant differences. For example, several of the protein synthesis inhibitors produced smaller sizes in *S*.*aureus* and larger in *E*.*coli* and *P*.*aeruginosa*.

**Figure 5.**
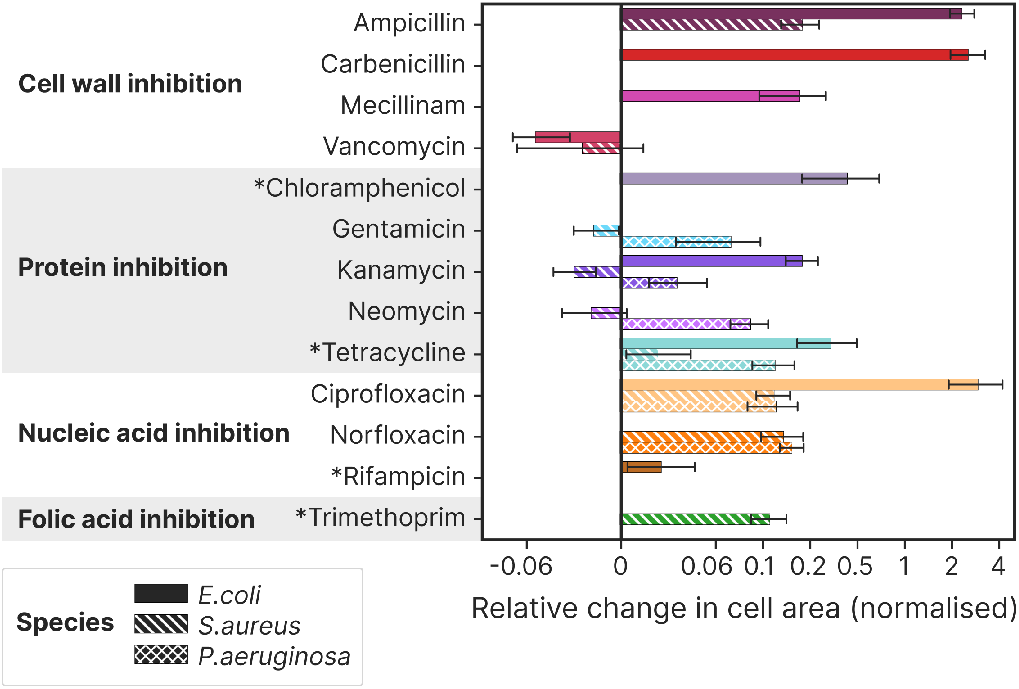
There are trends in morphology and growth characteristics between the antibiotic classes, but there are also exceptions. This plot shows how the mean cross-sectional area at the MIC is affected for the three species and for different antibiotics, with symlog x-scale. The antibiotics are grouped by action mechanism as inhibiting cell wall synthesis, protein synthesis, nucleic acid synthesis, and folic acid synthesis. The bacteriostatic antibiotics are marked with *, and the rest are generally considered bacteriocidal. Data is shown for *E*.*coli, S*.*aureus*, and *P*.*aeruginosa*. The error bars indicate a 95% confidence interval. The cell wall synthesis inhibitors produce the largest change in area.

### 2.5 MOR50 accurately estimates the MIC using just cell morphology and drastically reduces imaging time

As shown in fig. 3, the largest morphological changes occur in specific concentration ranges, beyond which they tend to subside at least to a certain degree. This is likely because when drug concentrations are very high, the transition between growth and growth halt/death is quick and does not allow enough time for morphological changes that depend on growth (see fig. S4-S6). We now asked how these morphologically relevant ranges related to the MIC. For all of the antibiotics and species considered, morphological changes were closely correlated with growth inhibition fig. S20-S22.

Motivated by these consistent morphology/MIC correlations, we developed a simple metric to estimate the MIC exclusively from morphological information acquired at a single time point (2.5 hours). *Morphological change 50* (MOR50), defines the MIC as the lowest antibiotic concentration that induces a change in cell area that exceeds 50% of the maximum cell-area change. For example, fig. 6.A shows ciprofloxacin concentration vs cell area after 2.5 hours of incubation with the antibiotic. The largest magnitude change is positive, so the MOR50 threshold is placed halfway between the control area at 0 µg mL^*−*1^ and max area at 0.015 µg mL^*−*1^, and the cell MIC is determined as the concentration where the mean cell area curve intercepts this line. For vancomycin in fig. 6.B, there is an overall larger reduction in area. Again, we place a threshold halfway between the control area and the maximum area change (which is now below the control area), and the cell MIC is determined as the concentration where the mean cell area curve intercepts this line. This process is shown for all combinations of tested antibiotics and species in fig. S23-S25. By normalizing antibiotic concentration to the concentration where growth is inhibited by 50% (*IC*_50_), and normalizing cell area to range between zero and one, fig. 6.C shows how well this works across the different antibiotics for *E*.*coli*. Similar plots for *S*.*aureus* and *P*.*aeruginosa* are shown in fig. S26.

**Figure 6.**
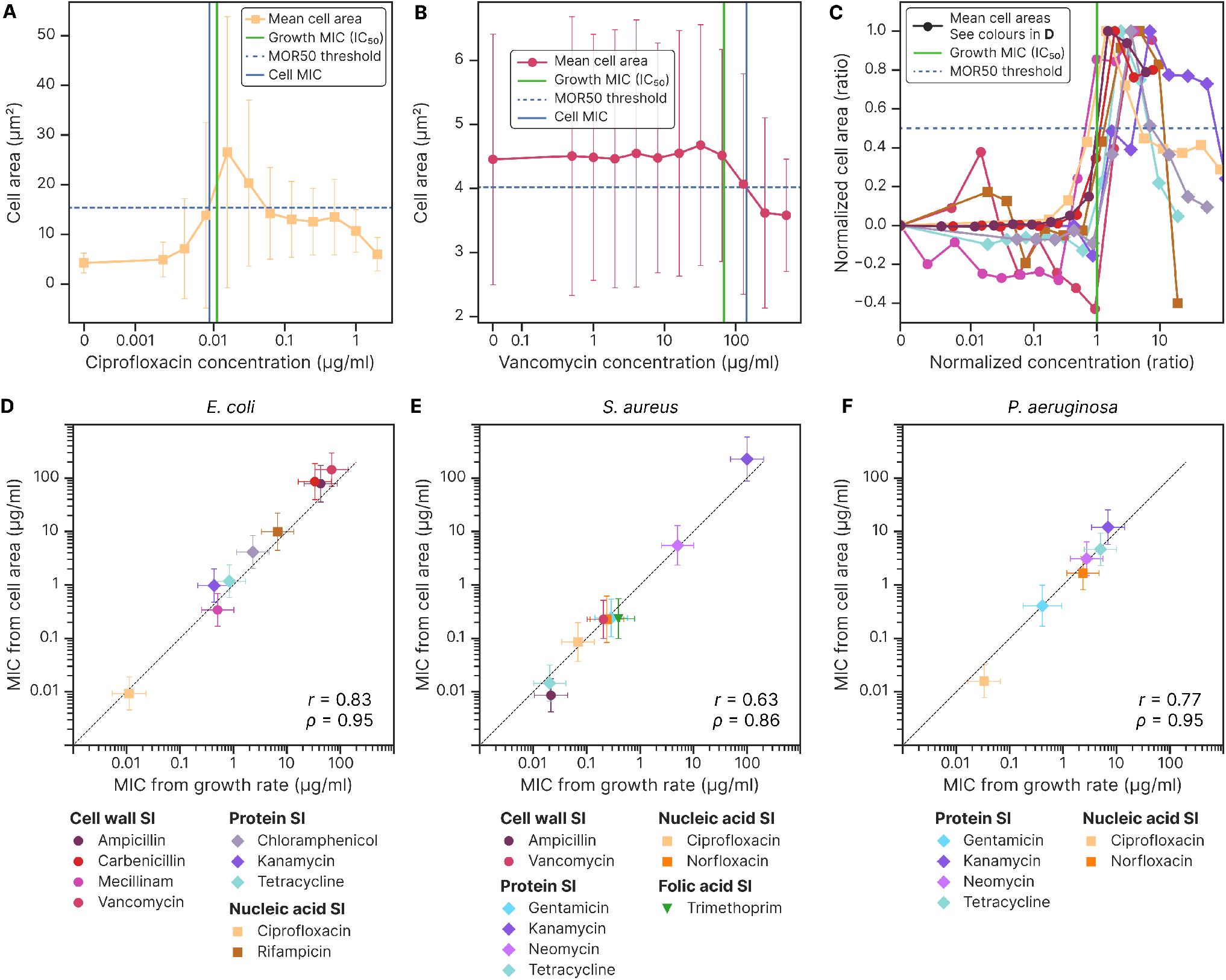
The magnitude of the morphology change is closely correlated with growth inhibition. MOR50 is a simple metric that can be used to determine the MIC from single-cell morphology data captured 2.5 hours after incubation with the antibiotics. **A** The mean cross-sectional cell areas for *E*.*coli* when exposed to varying concentrations of ciprofloxacin is shown (based on four repeats). The error bars show the standard deviation in cell areas. The MOR50 threshold is halfway between the area with no antibiotic present and the maximum area change and is shown as a dashed horizontal line. The cell MIC is where the area curve first crosses this threshold, represented by the blue vertical line. This corresponds very closely with MIC as determined by the growth rate, represented by the green vertical line. **B** For an antibiotic like vancomycin, the largest change is a reduction in area (*E*.*coli*, based on four repeats). The cell MIC is, therefore, determined to be where the area curve first crosses the MOR50 threshold from above. This corresponds well with the MIC as determined by the growth rate. **C** Showing how cell area changes depend on antibiotic concentration for *E*.*coli* and all the tested antibiotics. The antibiotic concentrations are normalized to growth MIC, and the cell area is normalized so that the no-antibiotic area is 0 and the maximum change in cell area is 1 (which for vancomycin entails inverting the area). The MOR50 threshold is shown as a horizontal line at 50% change, and the cell MIC is where the area curve first crosses this threshold. All the antibiotics produce a morphology change around the growth MIC. The degree of change varies strongly between antibiotics, which is here made apparent through the noise present in the area signal for antibiotics that induces a small change in morphology. The antibiotics are categorised by action mechanism as inhibiting cell wall synthesis, protein synthesis, and nucleic acid synthesis in addition to a negative control where no antibiotic was used, with legends displayed in subfigure **D. D** Comparing MIC from growth rate and cell morphology for *E*.*coli* shows a high degree of correlation, with annotated Pearson correction coefficient (*r*) and Spearman’s rank correlation coefficient (*ρ*). The dashed line is plotted for *y* = *x*. **E, F** The metric also generalizes to *S*.*aureus* and *P*.*aeruginosa*, each showing a similarly strong correlation between MIC as determined by growth rate and MIC as determined by cell morphology.

Comparing the MICs thus extracted with the MICs obtained from growth information alone, we observed a precise correspondence across the three microbial species fig. 6.D, E and F. A tabulated summary of the data is shown in table 1. The small amount of information required for the MOR50 makes it a powerful tool for MIC estimation as it minimizes imaging time. Using MOR50 and the MAP platform, we are able to determine the MIC of 8 antibiotics for a given biological sample in as little as 8 minutes of imaging after 2.5 hours of incubation when seeding from bacteria in the exponential growth phase.

**Table 1.** Showing average MIC values obtained through growth rate as outlined in [32] (Growth MIC), and single-cell morphology using the MOR50 metric (Cell MIC). All MIC values are reported in µg mL^*−*1^. Values are reported as mean and standard deviation between biological repeats. The combinations of species and antibiotics marked with “-” were not tested. We note that our strain of *S*.*aureus* carries a plasmid encoding for Kanamcycin resistance.

| Antibiotic | *E.coli* |  | *S.aureus* |  | *P.aeruginosa* |  |
| --- | --- | --- | --- | --- | --- | --- |
|  | Growth MIC | Cell MIC | Growth MIC | Cell MIC | Growth MIC | Cell MIC |
| Ampicillin | 70 ± 20 | 100 ± 60 | 0.021 ± 0.008 | 0.009 ± 0.003 | - | - |
| Carbenicillin | 50 ± 10 | 90 ± 60 | - | - | - | - |
| Chloramphenicol | 5.1 ± 0.8 | 4 ± 1 | - | - | - | - |
| Ciprofloxacin | 0.02 ± 0.01 | 0.009 ± 0.003 | 0.07 ± 0.02 | 0.09 ± 0.09 | 0.033 ± 0.007 | 0.016 ± 0.004 |
| Gentamicin | - | - | 0.29 ± 0.06 | 0.3 ± 0.1 | 0.4 ± 0.4 | 0.4 ± 0.4 |
| Kanamycin | 0.7 ± 0.3 | 1.0 ± 0.4 | 100 ± 30 | 200 ± 300 | 7 ± 2 | 12 ± 6 |
| Mecillinam | 1.0 ± 0.4 | 0.41 ± 0.04 | - | - | - | - |
| Neomycin | - | - | 5.07 ± 0.08 | 6 ± 7 | 2.8 ± 0.5 | 3 ± 1 |
| Norfloxacin | - | - | 0.2 ± 0.1 | 0.2 ± 0.4 | 2.4 ± 0.3 | 1.8 ± 0.5 |
| Rifampicin | 11 ± 2 | 10 ± 8 | - | - | - | - |
| Tetracycline | 2.0 ± 0.2 | 1.1 ± 0.3 | 0.020 ± 0.002 | 0.007 ± 0.008 | 5.0 ± 0.6 | 4.7 ± 0.7 |
| Trimethoprim | - | - | 0.39 ± 0.05 | 0.2 ± 0.2 | - | - |
| Vancomycin | 119 ± 5 | 140 ± 50 | 0.21 ± 0.03 | 0.2 ± 0.2 | - | - |

## 3 Discussion

The effects of antibiotics on the cell at a system level are only partly understood. Here, we used MAP, a high-throughput imaging platform we recently developed, to investigate the system-level effects of 13 antibiotics on the growth and morphology of three clinically relevant bacterial species. Specifically, we demonstrated that bacterial microcolonies grow exponentially at a certain, consistently heterogeneous, rate. Upon antibiotic treatment, as the dose approached the MIC, things changed. For the vast majority of the antibiotics we tested, different microcolonies responded heterogeneously to such drug doses such that their growth rates became markedly more heterogeneous. We observed the opposite trend for a small but self-coherent subset of antibiotics, the protein synthesis inhibitors, with a decrease in growth rate heterogeneity. Based on these observations, we speculate that bacteria’s growth rate heterogeneity in response to an antibiotic depends on the functional distance between antibiotic targets and cellular growth rate control. The active ribosomal pool has been recognised to modulate growth rate in a number of studies [37, 38, 39, 40, 41], and antibiotics that influence it directly benefit from a fast track towards growth impairment. Antibiotics that do not use this mechanism of action are functionally further away from growth rate control. Thus, their influence on growth rate is subject to the robustness of each single cell as a system. Cells are probably in slightly different states at the time of antibiotic treatment, but such differences are either of negligible entity towards the growth rate -the cells grow at the expected rate nevertheless -or they concern structures that are functionally distant or not directly involved in growth. Some examples of these could be repair mechanisms, or multidrug efflux pumps that only become relevant upon antibiotic exposure. It is tentative to hypothesise that such heterogeneity-boosting treatments may enhance persister production, for example, when drug cocktails are administered, motivating further studies. Lastly, the validity of these observations for *S*.*aureus* and *P*.*aeruginosa* further to *E*.*coli* suggests that, although these species have been studied less from the point of view of physiology at the system level, the same growth laws may hold true across the board.

Morphology is the other variable controlled at the system level that we were able to assay in this work. As demonstrated by others (primarily using data from the balanced growth of *E*.*coli*), many antibiotics alter bacterial morphology. Here, we have shown that this is true for all of the 24 antibiotic-bacterium combinations, independent from a cell’s rod or coccoid shape and antibiotic mechanism of action, even though this does play a role in determining the magnitude. We have shown that defects require growth to emerge in all cases, although we did not investigate antimicrobials such as Gramicidin S, that directly pierce the cell envelope [42]. We could confirm the well-established mechanism that leads to morphological alterations for some of the antibiotics tested, such as ciprofloxacin or betalactams. In other cases, the results were surprising. For example, in a rich medium such as the one in our pads, *E*.*coli* cells in balanced growth that are treated with chloramphenicol are expected to shrink [4] while we observed them swell. While we do not know the reason for this discrepancy, we assume that this emerges from the differences in growth conditions.

We have further shown that single-cell morphology alone can accurately estimate the minimum inhibitory concentration (MIC) of antibiotics, providing a rapid and cost-effective method for single-cell imaging research and antibiotic susceptibility testing (AST). *Morphological change 50* (MOR50), the morphology-based metric we introduced correlates well with traditional growth-rate-based MIC determination for all tested antibiotic classes, and can provide MIC estimation within 2.5 hours of incubation, with only a few minutes of imaging time. This approach has the potential to significantly improve AST by reducing the time and cost required for testing, thereby aiding in the selection of appropriate antibiotics for effective infection treatment and providing a new tool against the rise of antimicrobial resistance.

## 4 Materials and Methods

### 4.1 Sample preparation

Experiments were conducted with strains of *E*.*coli, P*.*aeruginosa* and *S*.*aureus*. See table 2 for details about the species and growth medium used. We note that while the *S*.*aureus* strain we used was resistant to Kanamycin, as shown in fig. S27, the gentamycin resistance was excised from our PA01-CFP strain as confirmed by PCR. All of the bacteria morphology data presented in this work was collected using the Multipad Agarose Platform (MAP) as described in [32]. The dataset for *E*.*coli* used in this study is the same as that in our previous work [32], where the MAP platforms and the initial sample preparation methods were identical. The MAP platforms were prepared in batches of three, with antibiotic dilution series as outlined in fig. S1. Pre-cultures were grown overnight, diluted 500x into fresh media, and allowed to resume exponential growth for at least three generations. These seed cultures were then diluted to a 600 nm optical density (OD600) of 0.02, and aliquots of 1.5 µL were placed on the surface of each MAP pad. The pads were dried for 5 to 10 minutes before peeling the protective film on the upper adhesive sheet and sealing the pads with a single glass slide that covers all the pads (UQG Optics, GPD-1577, dimensions 110 *×* 74 *×* 0.17 mm).

**Table 2.** Outlining the strain, growth media and analysis statistics for the different bacteria species used for these experiments. The growth media was Luria-Bertani (LB) Broth (ThermoFisher, 10855001, with 10 g peptone, 5 g yeast extract, 5 g sodium chloride per 1 L media) and Mueller Hinton (MH) Broth (Sigma-Aldrich, 70192, with 2.0 g beef infusion solids, 17.5 g casein hydrolysate and 1.5 g starch per 1 L of media). We started using LB Broth with *E*.*coli*, but switched to MH Broth for the later experiments with *P*.*aeruginosa* and *S*.*aureus* as we wanted to test with this media as well, given it is recommended for use with AST by EUCAST [43]. The same growth media was used for precultures and preparing pads on the MAP platform. Pad count, microcolony count and bacteria count show the total numbers included in the datasets used in this work. These numbers are reported from a single timepoint around 2.5 hours after imaging was started, meaning each microcolony and cell counted are unique. For the analysis, data from a few consecutive time steps is typically used.

| Species | Strain | Growth media | Pad count | Microcolony count | Bacteria count |
| --- | --- | --- | --- | --- | --- |
| *E.coli* | K-12, MG1655 | LB Broth | 437 | 4 155 | 388 576 |
| *P.aeruginosa* | PA01-CFP [44] | MH Broth | 395 | 5 796 | 234 460 |
| *S.aureus* | SH1000-pMV158GFP [45] | MH Broth | 447 | 4 887 | 171 411 |

### 4.2 Timelapse microscopy

The MAP platform is imaged at 37 ^*°*^C on a custom-built, open-frame inverted microscope for 5 hours. One field of view (FOV) is imaged per pad, using a looping script to capture all the images automatically. An LED focusing system is used to keep the sample roughly in focus automatically. To account for possible misalignment between the camera’s focal plane and the imaging plane of individual pads (which can exhibit significant tilts in relation to each other on the same MAP), a z-stack of images is captured for each FOV. We typically capture about ten frames per stack with 0.4 µm step size. This approach ensures that each microcolony is captured at its optimal focus in one of the frames, despite the spatial variations in the imaging planes across different pads. For these experiments, we chose the FOVs manually at the start of the time-lapse. However, selecting FOVs could be done in an automated fashion to give a fully automated imaging workflow. Our workflow enables the rapid high-throughput imaging of a large number of microcolonies and cells, their numbers for this study are given in table 2.

Imaging was performed with brightfield illumination using a Nikon 40x CFI Plain Flour air objective with a numerical aperture of 0.75. The camera was a Teledyne FLIR BFS-U3-70S7M-C with a 7.1 MP Sony IMX428 monochrome image sensor. All images were captured at 3208 *×* 2200 pixels, resulting in an effective resolution of 0.112 µm*/*pixel. The temporal resolution of the datasets is about 8 minutes, limited by the time it takes for the microscope to image all 96 pads in the MAP.

### 4.3 Antibiotics

The antibiotics were mixed into separate dilution series before being transferred to the pads of the MAP. They were sourced as follows: ampicillin (Sigma-Aldrich 10835242001), carbenicillin (Sigma-Aldrich C1389), ciprofloxacin (Sigma-Aldrich 17850), chloramphenicol (Sigma-Aldrich C0378), gentamycin (Sigma-Aldrich G1397), kanamycin (Sigma-Aldrich K4000), mecillinam (Sigma-Aldrich 33447), neomycin (Sigma-Aldrich N1142), norfloxacin (Sigma-Aldrich N1142), tetracycline (Sigma-Aldrich T3258), trimethoprim (Sigma-Aldrich T7883), rifampicin (Sigma-Aldrich 557303) and vancomycin (Sigma-Aldrich V2002). Antibiotics were homogeneously blended with the agarose and growth medium at 60 ^*°*^C prior to preparing the dilution series. Subsequently, the mixture was dispensed onto the MAP platform using an Opentrons OT2 pipetting robot [46], where it solidified to form the small assay pads. Stock solutions were prepared by dissolving the antibiotics in 10 mL of milliQ water (ampicillin, carbeni-cillin, ciprofloxacin, gentamycin, kanamycin, mecillinam, neomycin, tetracycline, vancomycin), 100% methanol (rifampicin, trimethoprim), or 95% ethanol (chloramphenicol, norfloxacin). These were stored at *−*20 ^*°*^C until use. Some antibiotics were subjected to ultrasonication to aid in their dissolution process.

### 4.4 Image processing

The images are analysed using the processing pipeline available in *PadAnalyser*. The process of z-stack projection and colony segmentation is outlined in [32]. In addition, single-cell segmentation is developed and demonstrated here, performed in a series of steps outlined in fig. S2. We start by computing the laplacian of the Gaussian of the brightfield image and then using a simple threshold to binarize the image. The binary image is filtered using the colony masks, and converted to contours with OpenCV [47]. Very small contours are removed as they correspond to debris and optical artefacts, and the remaining contours are split based on an outline curvature metric and point separation to make sure neighbouring bacteria get individual masks fig. S3. Finally, the contours are dilated to represent the true cell areas better. The colony and single-cell statistics are bundled into two data frames per experiment: one for the colony statistics (with single-cell statistics linked to each relevant colony) and one for the single-cell statistics. Similarly, two output video files are produced for each field of view with segmentation masks for colony outlines and single cells drawn in clear colours so the user can validate that the algorithms are working correctly, see [48].

We could not find one set of parameters that would accurately segment the three different species, so we determined four key segmentation parameters we could alter to tune the algorithm on a species-by-species basis. These are outlined in tab. S1 along with the values used. These parameters are chosen to ensure the segmentation pipeline is robust to the changes in cell morphology observed for each species.

The single-cell segmentation code was compared to manually annotated images to ensure accuracy. We manually labelled frames from a range of different time points and antibiotic concentrations for all species to ensure the approach is robust to different cell morphologies, colony sizes and imaging conditions. The comparison results are outlined in tab. S2, highlighting three main metrics. *Intersection over union* (IoU) outlining the overall area overlap, *mean cell area error* as a ratio between the mean cell areas, and *mean errors per cell* report the number of segmentation errors per cell in the labelled image [49].

### 4.5 Extracting single cell statistics

In *PadAnalyser*, we compute the area (*A*) of rod-shaped bacteria using the *cv2*.*contourArea* function from the OpenCV library [47], which accurately calculates the area of the mask representing the bacteria. For width estimation, we utilized the Euclidean distance transform (EDT), taking twice the maximum value of the EDT as the representative width (*w*) of the bacteria. This approach effectively captures the diameter of the widest part of the bacteria. The length (*l*) was then computed using the formula 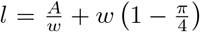 This formula derives from the relationship between the area, width, and a correction factor accounting for the curvature and shape irregularities of the bacteria. The term 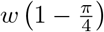 serves as an adjustment to the basic area-to-width ratio, compensating for the fact that the bacteria are not perfect rectangles, thus providing a more accurate measure of length for rod-shaped and potentially curved bacteria.

## Supplementary Information

Supplementary information is available for this paper containing additional data and figures that provide further insight into the study.

## Acknowledgments

We thank Erika Causa, Viridiana Carmona Sosa, Aske Petersen, and Patryk Nienaltowski for insightful feedback and support. The project was funded by the EU EC ITN Phymot, Marie Sklodowska-Curie grant agreement No 955910. LM acknowledges funding from the Herchel Smith Postdoctoral Fellowship.

## Data availability statement

Our open-source Python package, PadAnalyser, can be found at github.com/Cicuta-Group/PadAnalyser and installed through the PyPI package manager [50]. This package contains the entire analysis pipeline we use for image preprocessing, segmentation, statistics extraction and plotting. Instructions for how to make, assemble and use the MAP platform can be found at github.com/Cicuta-Group/MAP-imaging [51], along with an example dataset and a demo showing how to use the PadAnalyser package. The data from experiments used in this work can be found at doi.org/10.5281/zenodo.10352996 [48].

## Declarations

AD declares to work for a company that operates in the application areas described in this work, but has no direct conflict of interest.

## Notes

https://github.com/Cicuta-Group/PadAnalyser

https://github.com/Cicuta-Group/MAP-imaging

https://doi.org/10.5281/zenodo.10352996

