## Supplementary information for "Antibiotics Change the Growth Rate Heterogeneity and Morphology of Bacteria"

### Supplementary Information for
### Antibiotics Change the Growth Rate Heterogeneity and Morphology of Bacteria

Morten Kals[a,b], Emma Kals[a], Jurij Kotar[a], Allen Donald[b], Leonardo Mancini*[a,1], and Pietro Cicuta*[a,2]

[a]Cavendish Laboratory, University of Cambridge, Cambridge CB3 0HE, UK
[b]Synoptics Ltd., Cambridge CB4 1TF, UK
[1]
[2]
*joint last authors

#### This PDF file includes

- Tables S1 to S3.
- Figures S1 to S27.

Table S1: An overview of the parameters used to tailor the segmentation pipeline to the different species of bacteria. Sigma sets the strength of the Gaussian blur used to smooth the initial z-stack projected image before computing the Laplacian. The min mask size filter sets the minimum size of a mask to be considered a cell. Threshold sets the threshold value used to binarise the image. The split factor controls how aggressively the masks are split.

| Parameter | *E. coli* | *S. aureus* | *P. aeruginosa* |
|---|---|---|---|
| sigma | 1.5 | 2.5 | 1 |
| threshold | -1000 | -2000 | -3000 |
| min mask size filter (px) | 60 | 15 | 30 |
| split factor | 0.3 | 0.75 | 0.65 |

Table S2: Comparison of how accurate the PadAnalyser segmentation code is based on the metrics *intersection over union* (IoU), *mean cell area error*, and *mean errors per cell* across different species. The data includes total frame count and cell count to indicate the extent of the comparison. The row *manually annotated* compares two frames that both have been manually annotated to assess best-case metrics.

| Species | Frame count | Cell count | IoU | Mean cell area error | Mean errors per cell |
|---|---|---|---|---|---|
| *E. coli* | 10 | 2421 | 0.76 | 0.11 | 0.39 |
| *S. aureus* | 7 | 642 | 0.77 | 0.19 | 0.21 |
| *P. aeruginosa* | 7 | 2010 | 0.75 | 0.039 | 0.44 |
| Manually annotated | 3 | 369 | 0.77 | 0.0099 | 0.13 |

Table S3: EUCAST MIC concentrations are based on tabulated confidence intervals from mic.eucast.org (see table S3). Where data is available but the confidence interval is not reported, the mean and standard deviation of available data are used to estimate the confidence interval.

| Antibiotic | *E. coli* | *S. aureus* | *P. aeruginosa* |
|---|---|---|---|
| Ampicillin | $10 \pm 6$ | $0.3 \pm 0.2$ | - |
| Carbenicillin | - | - | - |
| Chloramphenicol | $12 \pm 4$ | $12 \pm 4$ | - |
| Ciprofloxacin | $0.05 \pm 0.02$ | $1.5 \pm 0.5$ | $0.8 \pm 0.3$ |
| Gentamycin | - | - | - |
| Kanamycin | $20 \pm 10$ | $30 \pm 30$ | $500 \pm 500$ |
| Mecillinam | $0.4 \pm 0.1$ | - | - |
| Neomycin | $5 \pm 4$ | $2 \pm 2$ | $100 \pm 100$ |
| Norfloxacin | $0.3 \pm 0.2$ | $8 \pm 8$ | $4 \pm 4$ |
| Rifampicin | - | $0.012 \pm 0.004$ | - |
| Tetracycline | $3 \pm 1$ | $0.6 \pm 0.4$ | $80 \pm 50$ |
| Trimethoprim | $3 \pm 2$ | $1.3 \pm 0.8$ | - |
| Vancomycin | - | $1.5 \pm 0.5$ | - |

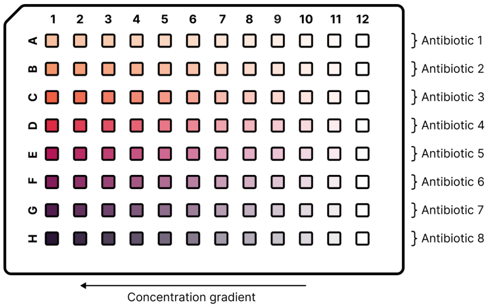


Figure S1: MAP setup used for AST where each row has a concentration gradient of a different antibiotic, making it possible to test eleven concentrations + control for eight combinations of antibiotic and bacteria at a time.

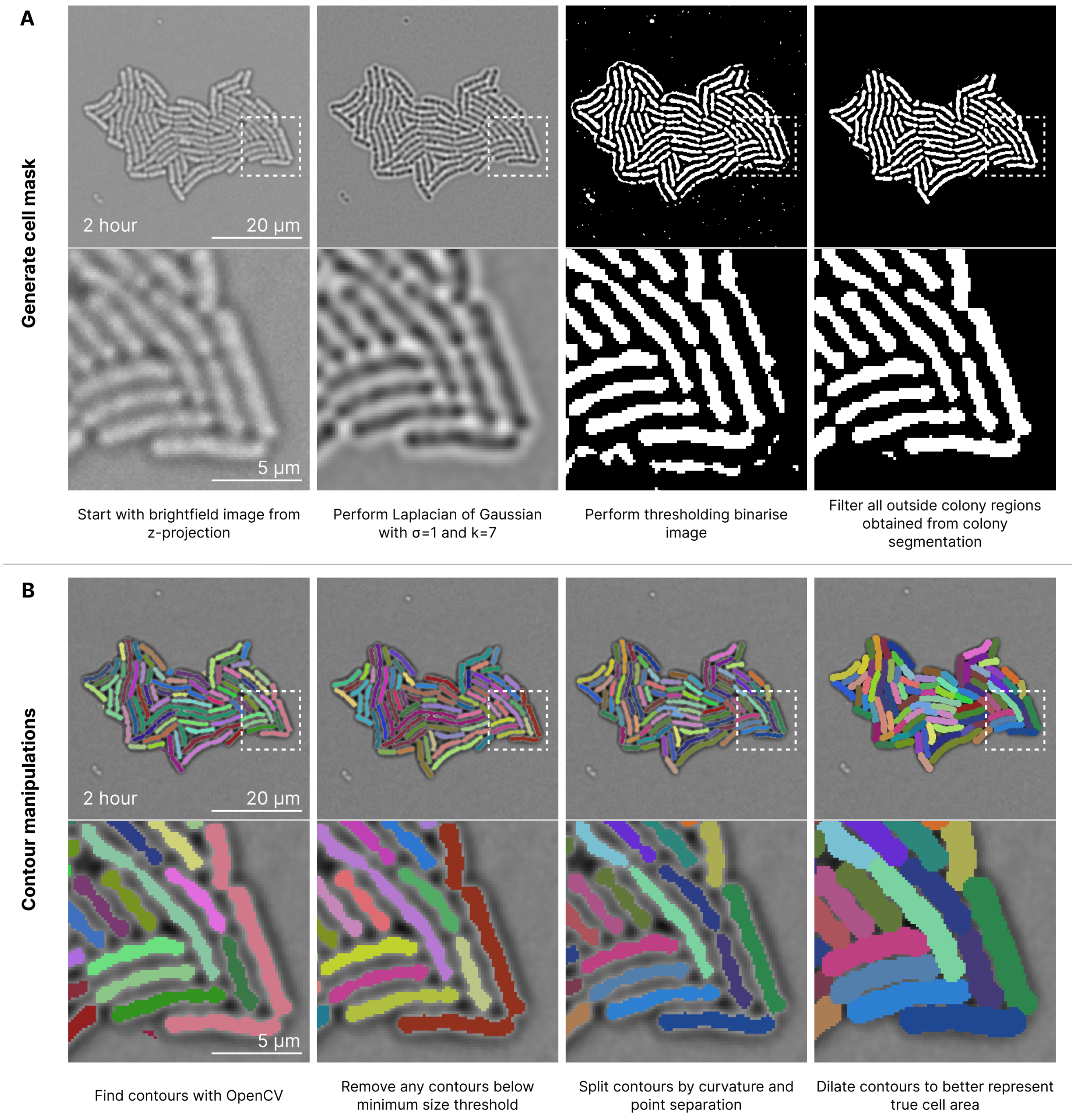


Figure S2: Step-by-step view of the cell segmentation process. A sample colony is used to illustrate each step, with a zoomed-in view for each step to highlight the cell details. The colony was captured after two hours of imaging. **A** First, the z-stack projection of the brightfield image is processed to generate a binary image containing the cell masks. **B** Then, the masks are converted to contours and manipulated to produce accurate segmentation for the bacteria.

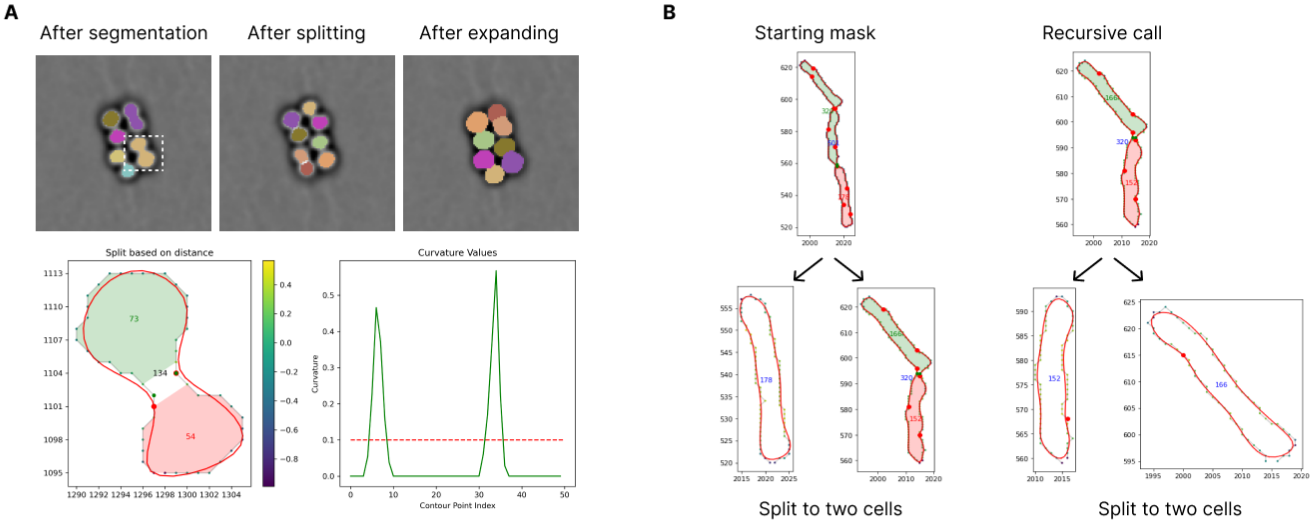


Figure S3: An overview of how the contours are split into separate cells based on outline curvature and separation. **A** shows an example with *S. aureus*. A spline is fitted to the outline of the cells (shown in red), and its curvature is computed at each point. Positive curvature maps to where the surface curves outwards. Discarding all negative curvatures, the maximum curvature locations are found. If more than 10 points separate their position along the curve, and they are closer than the max width of the contour times the split factor, the split is performed. Using these points of maximum curvature as seed points, a local search is conducted around these points to find the pair of points that are closest to each other. **B** shows an example with *E. coli* where the recursive nature of the algorithm is highlighted.

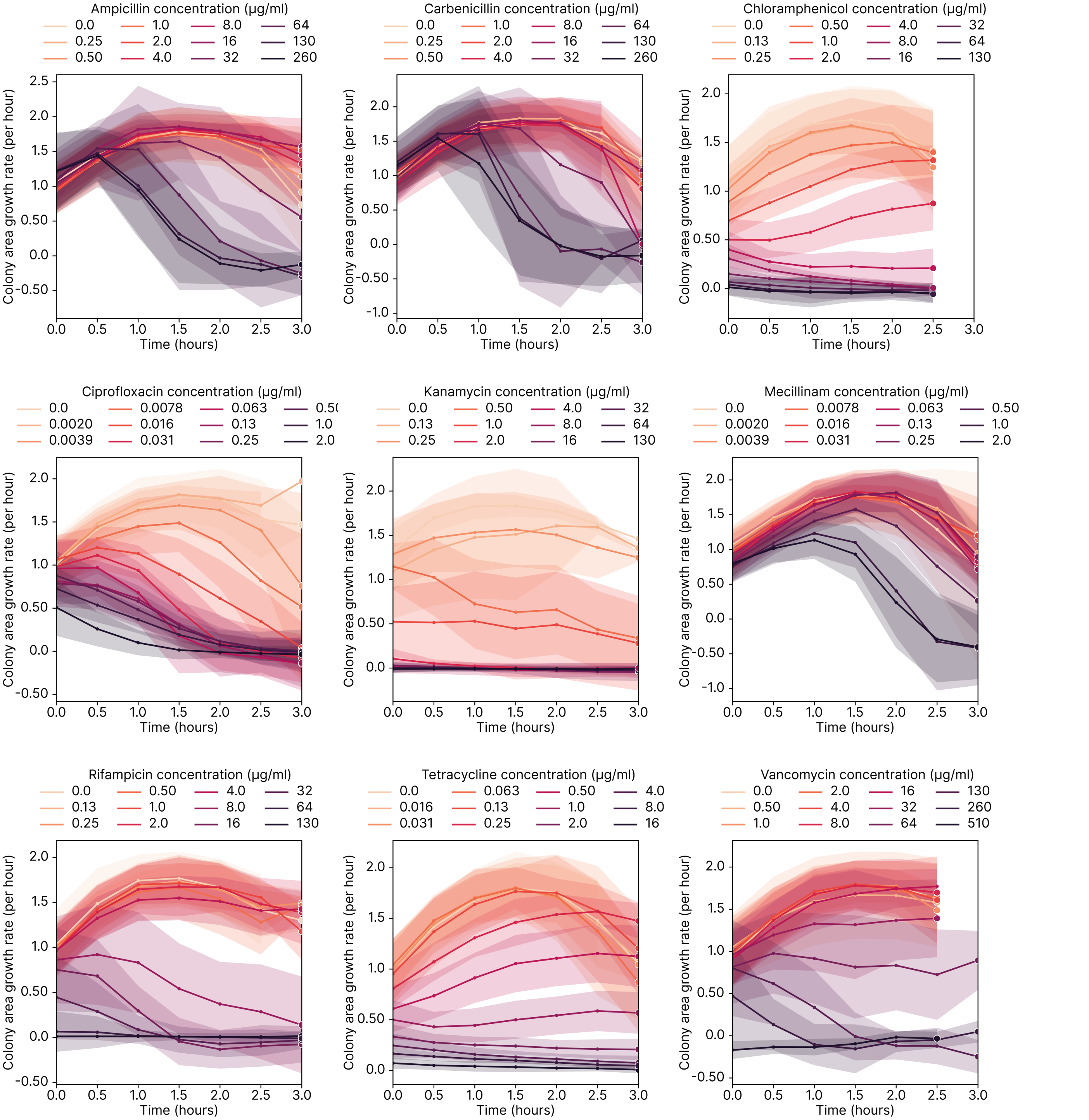


Figure S4: Overview of how the antibiotics impact the growth rate of *E. coli* over time. The growth rate is computed as the time derivative of the growth curve, and the shaded region represents the standard deviation of the growth rate. Darker lines represent higher antibiotic concentrations. This figure is based on the same *E. coli* dataset used in [Kals et al.(2024)Kals, Mancini, Kotar, Donald, and Cicuta], and is included here for completeness.

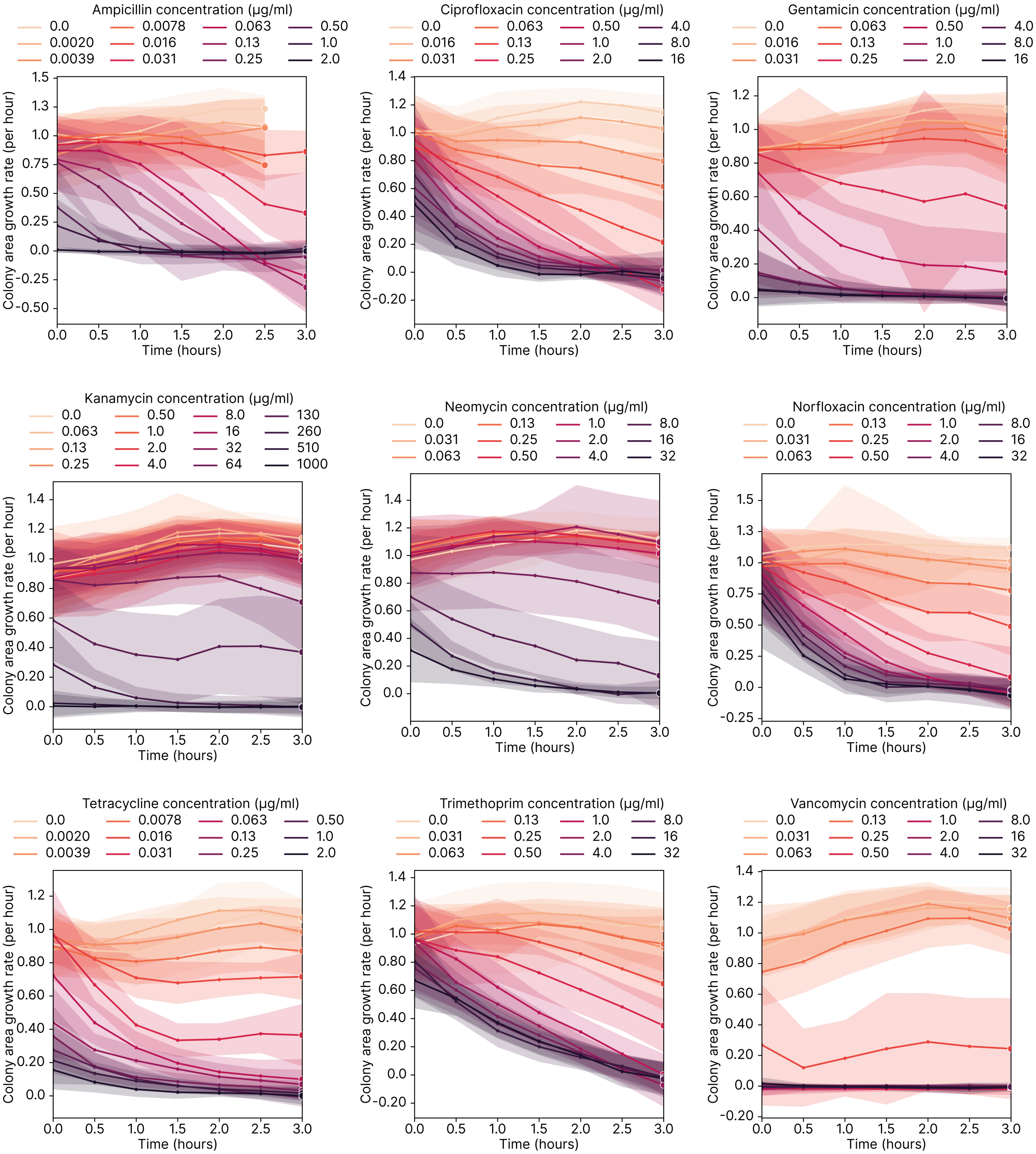


Figure S5: Overview of how the antibiotics impact the growth rate of *S. aureus*, see legend of fig. S4 for details.

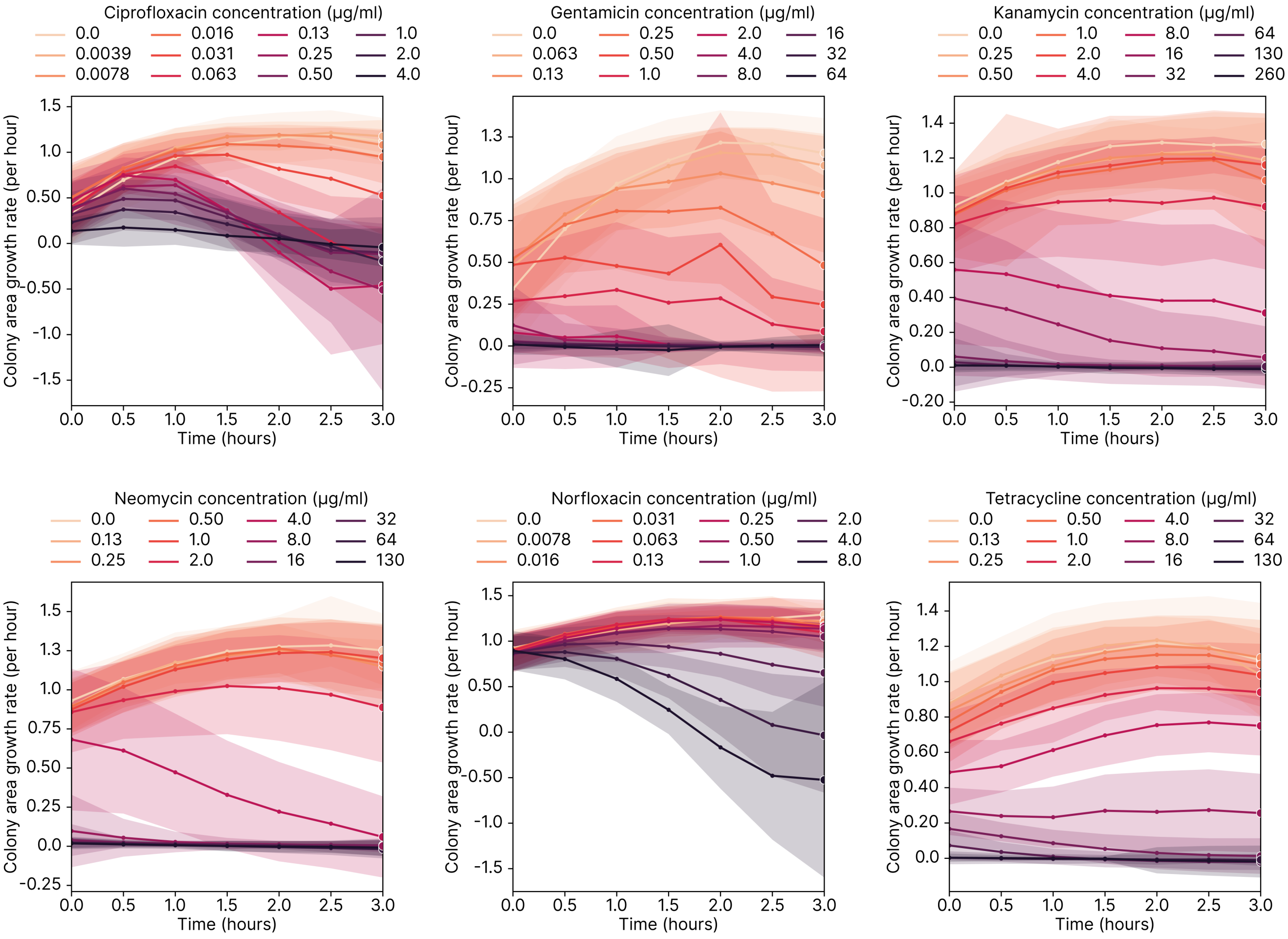


Figure S6: Overview of how the antibiotics impact the growth rate of *P. aeruginosa*, see legend of fig. S4 for details.

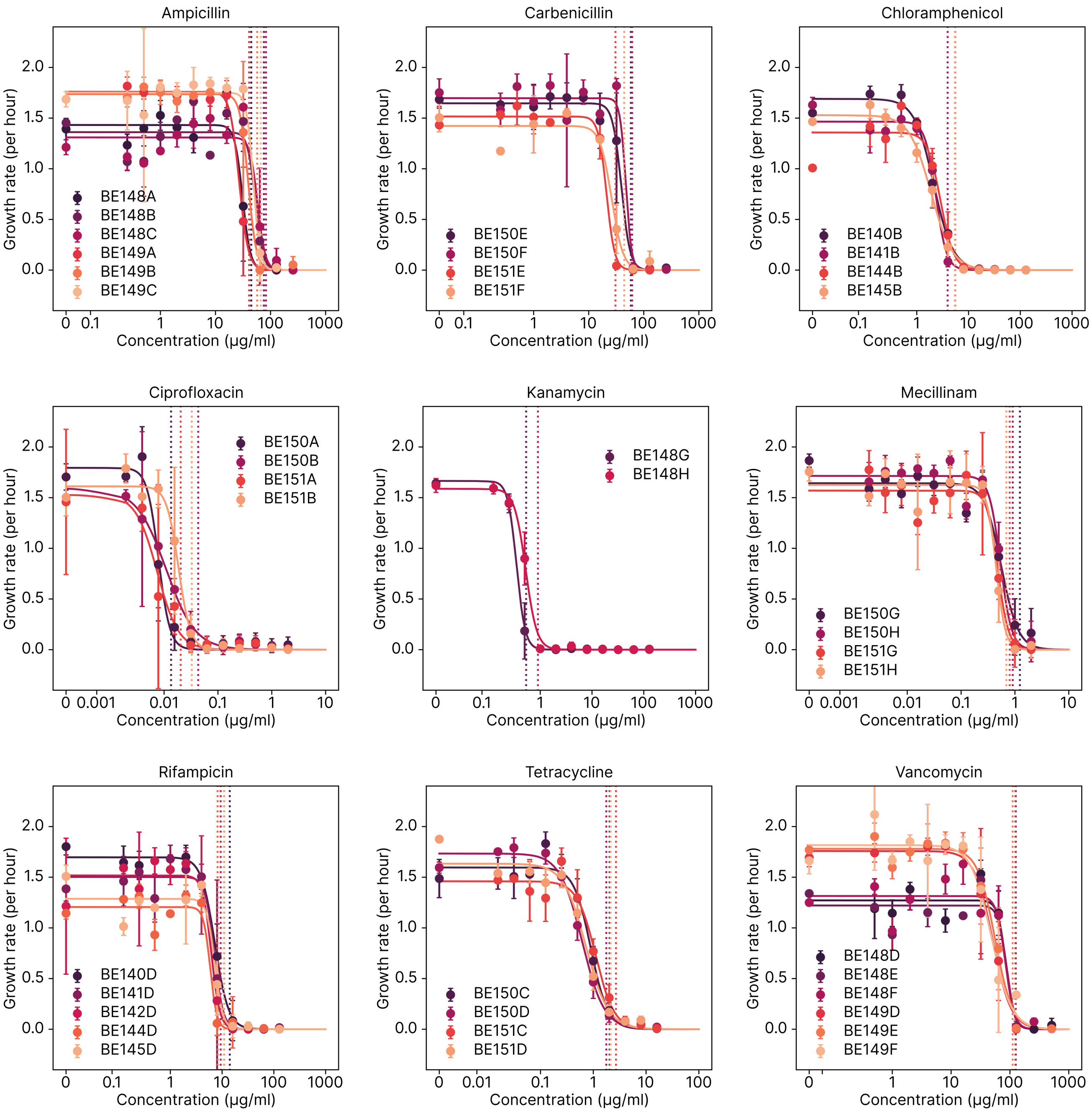


Figure S7: Growth rate dependence on antibiotic concentration for *E. coli*. Each repeat is plotted individually, and Hill fits performed for each repeat. Data used is for the time between 2 and 3 hours of growth. The vertical lines signify IC90 concentrations where growth is inhibited by 90%. This figure is based on the same *E. coli* dataset used in [Kals et al.(2024)Kals, Mancini, Kotar, Donald, and Cicuta], and is included here for completeness.

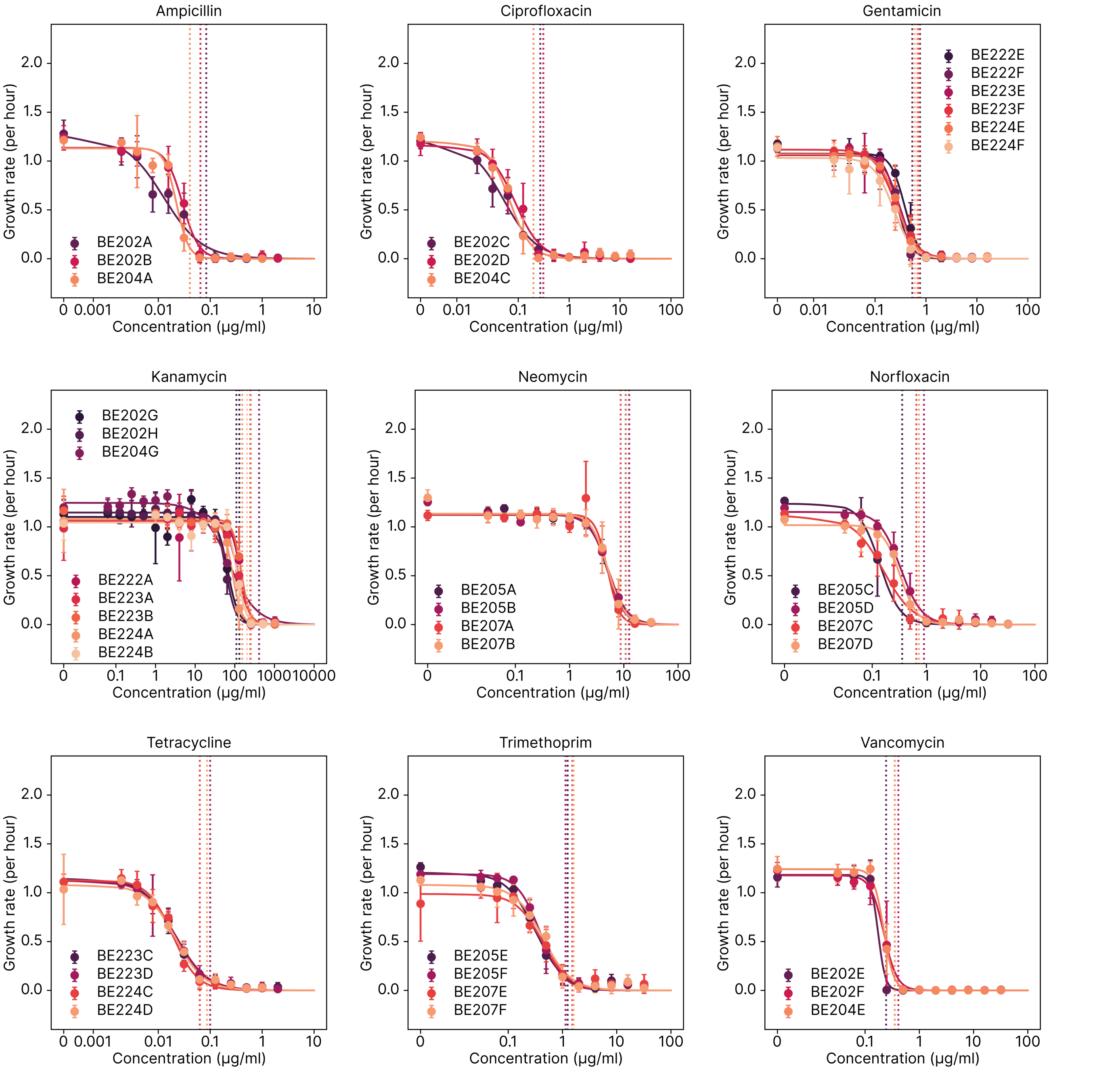


Figure S8: Growth rate dependence on antibiotic concentration for *S. aureus*, see legend of fig. S7 for details.

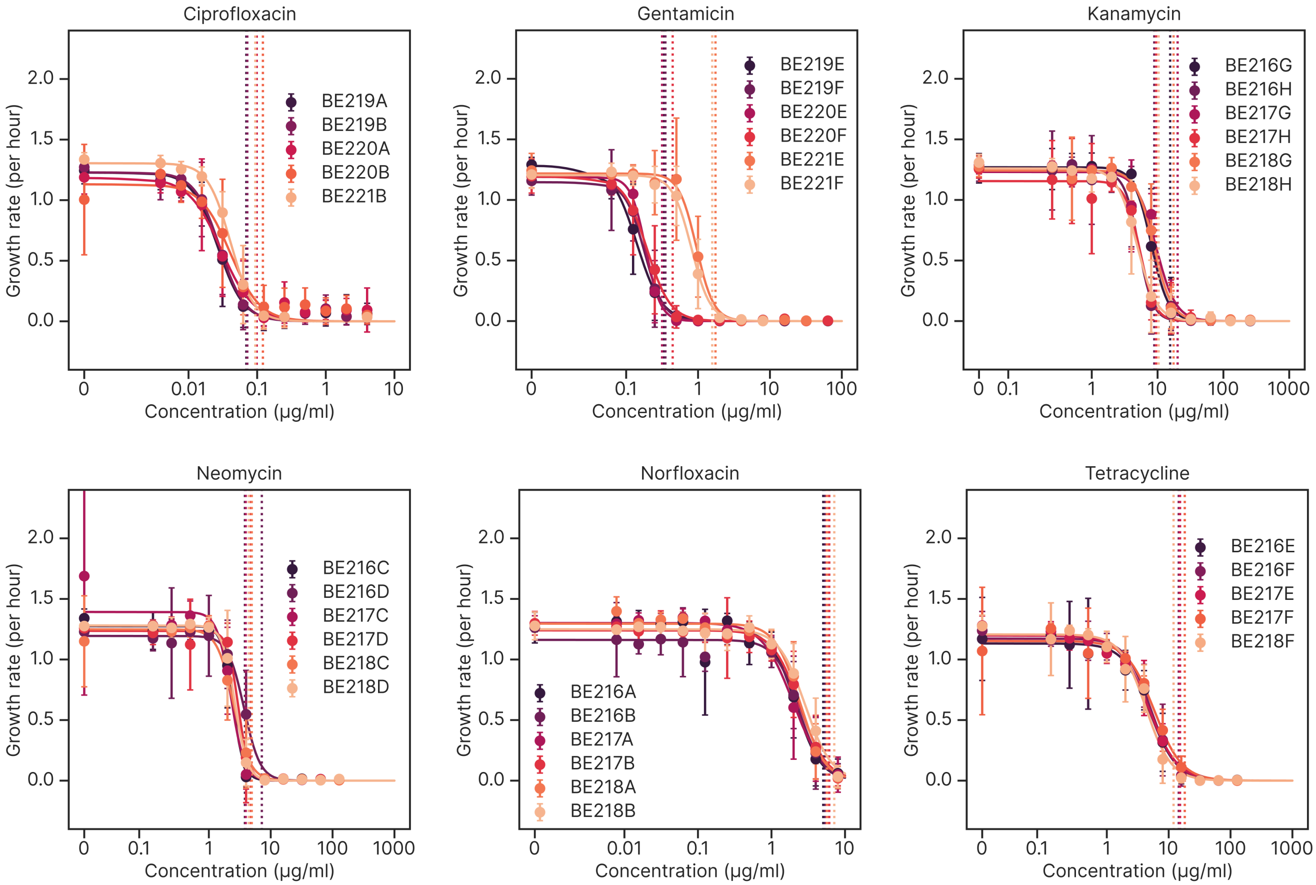


Figure S9: Growth rate dependence on antibiotic concentration for *P. aeruginosa*, see legend of fig. S7 for details.

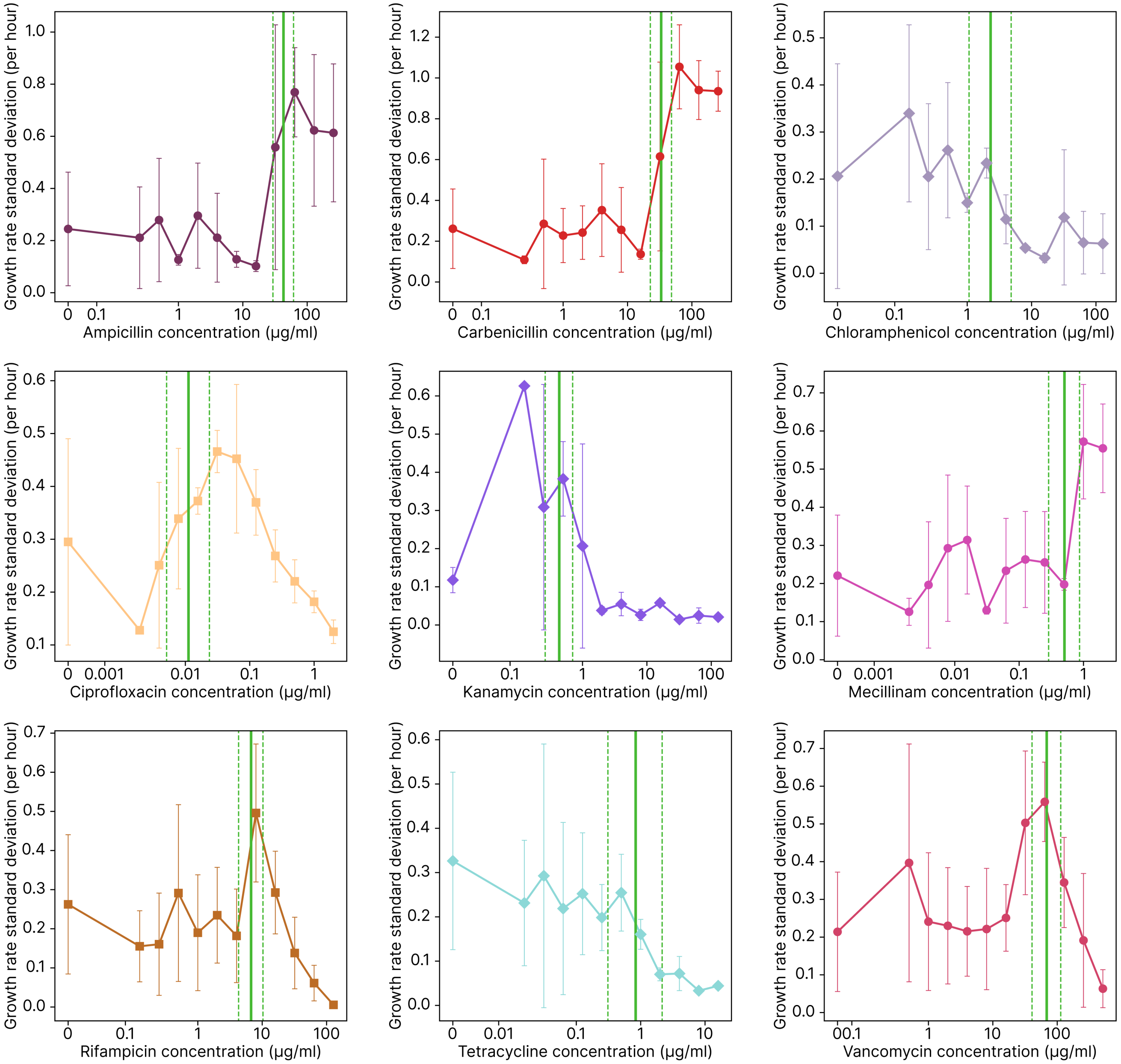


Figure S10: Showing the growth rate heterogeneity (GRH) at different concentrations for *E. coli*. The variation in growth rate on a single pad is computed and averaged across repeats in the 1 to 2.5-hour time period. The data is averaged across all repeats shown in previous graphs (two to eight per antibiotic/species combination). The vertical lines correspond to the MIC concentrations, with $IC_{10}$ at the lowest concentration, then $IC_{50}$ and $IC_{90}$ at the highest concentration. We see that for some antibiotics, high concentrations produce a much higher variation in growth rate between the colonies and that this change is linked to the MIC. For others, like tetracycline, there is a drop in GRH linked to the use of the antibiotic, also around the MIC.

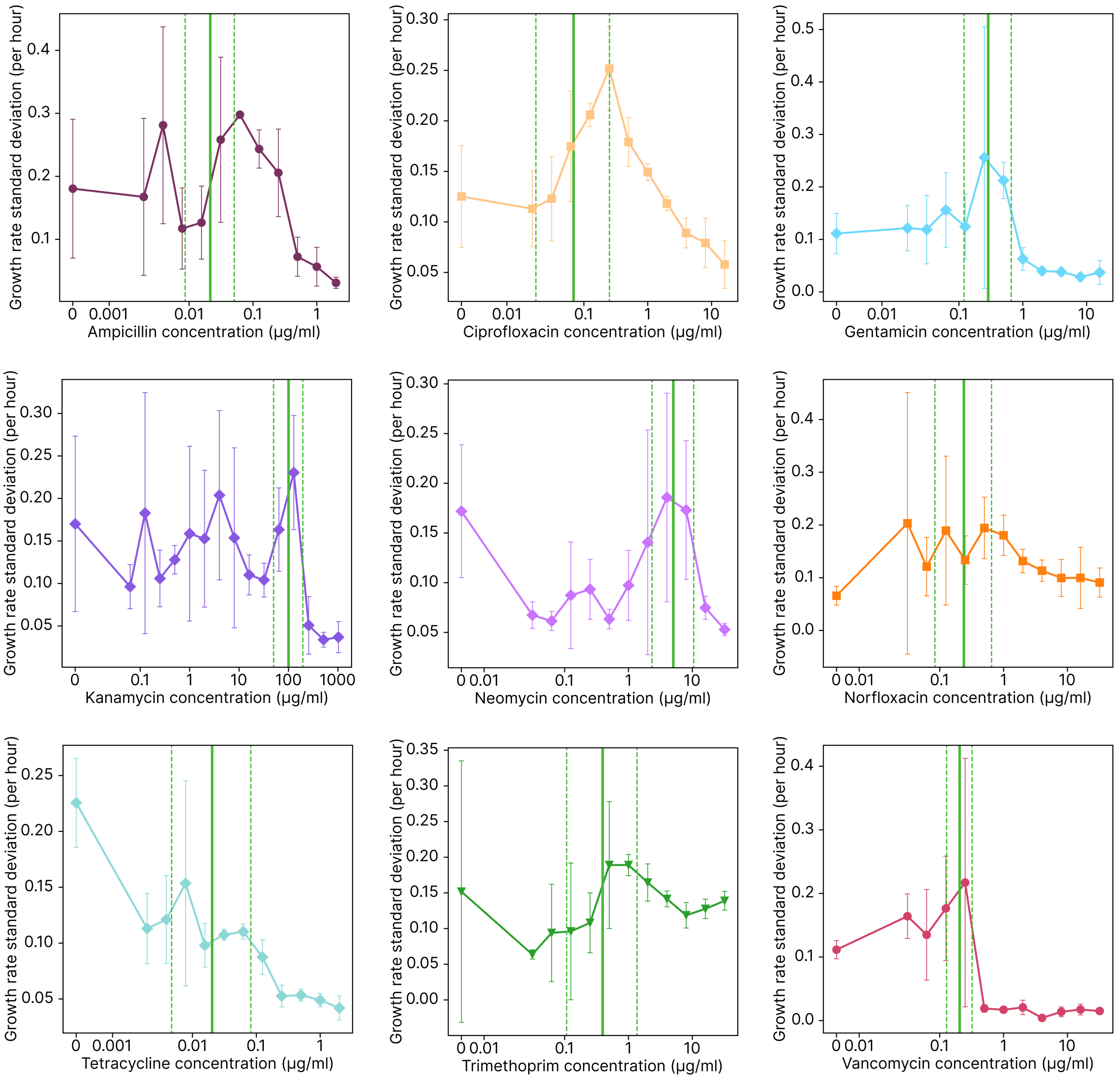


Figure S11: Showing the GRH at different concentrations for *S. aureus*, see legend of fig. S10 for details.

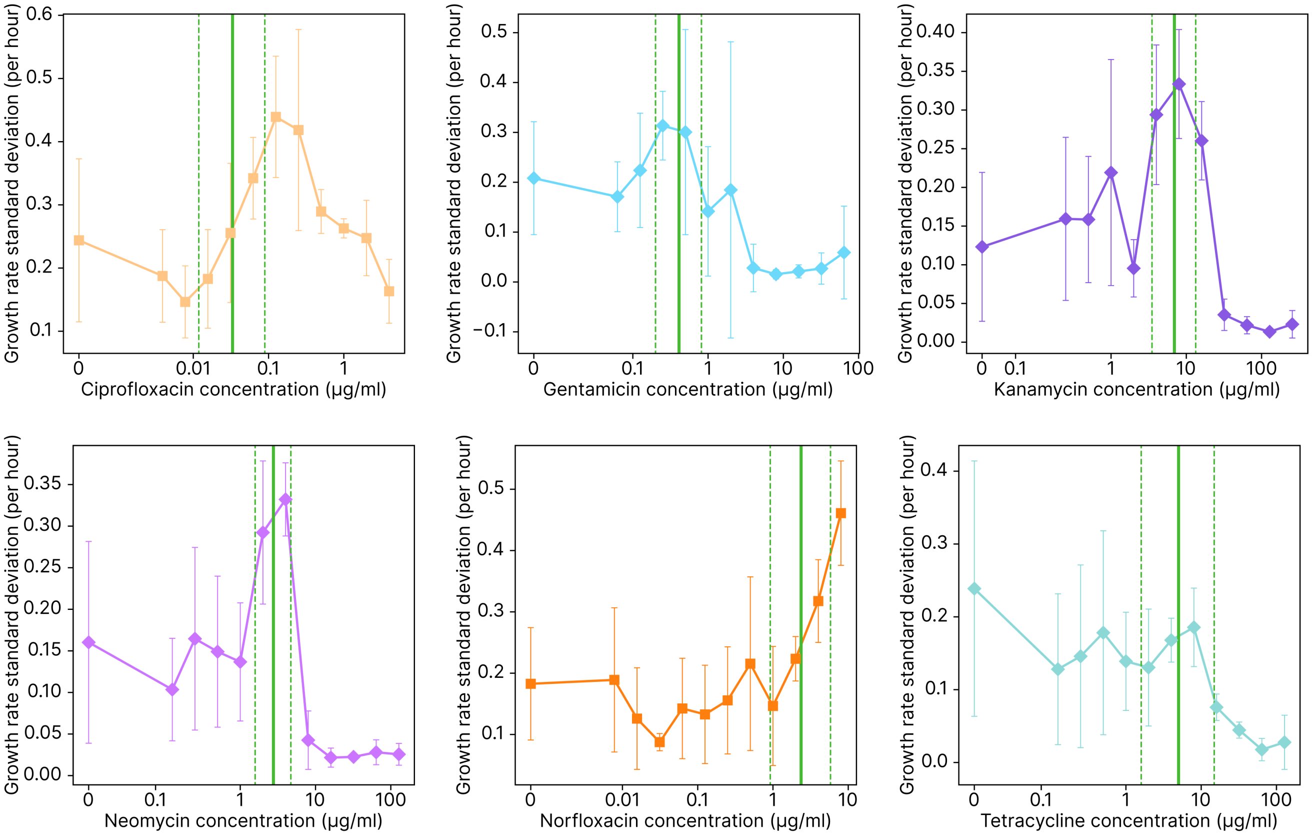


Figure S12: Showing the GRH at different concentrations for *P. aeruginosa*, see legend of fig. S10 for details.

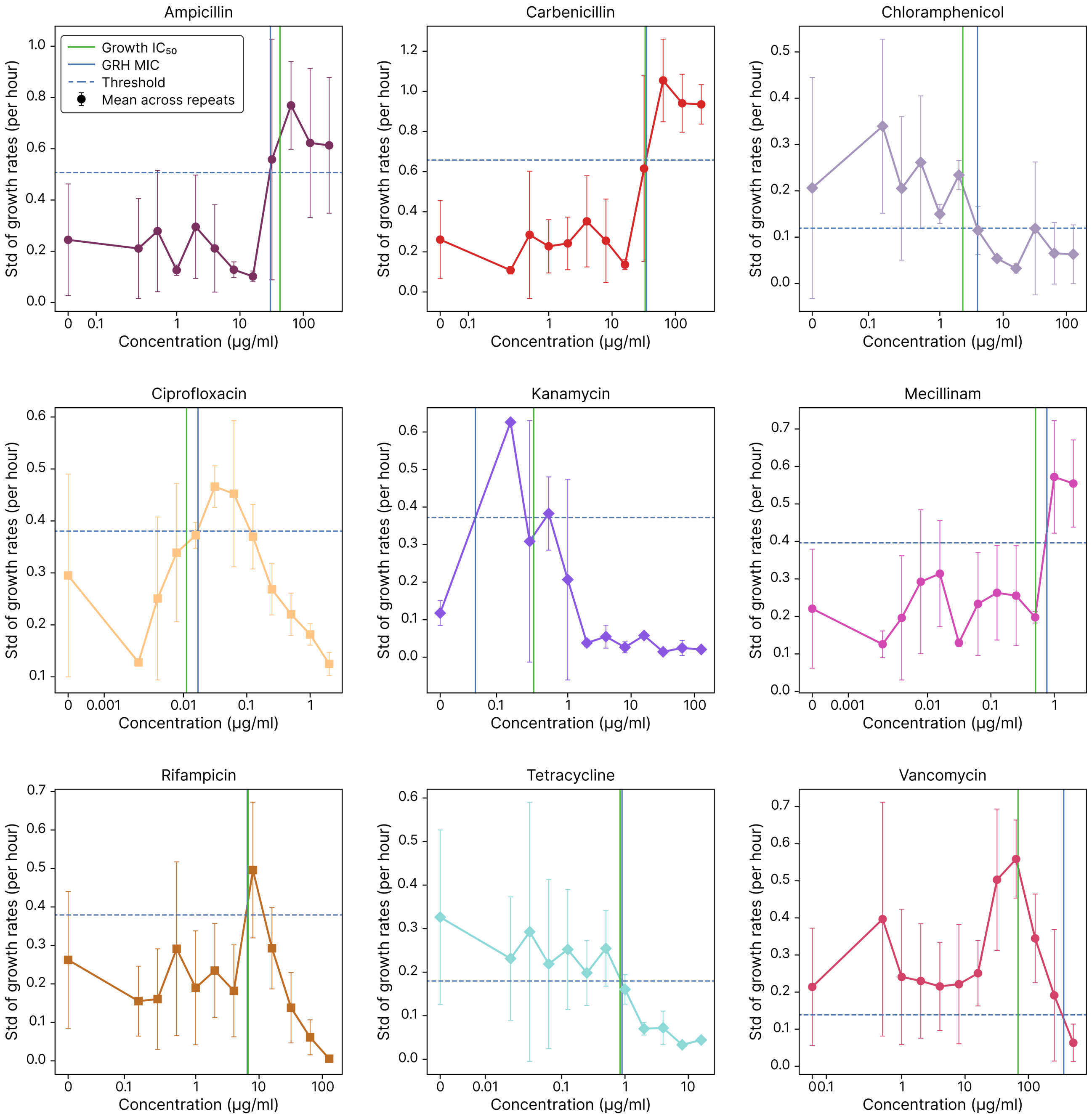


Figure S13: Showing how GRH can be used as a metric to determine MIC for *E. coli*. The data is the same as used in fig. S10, but here we can determine MIC based on this signal by applying a dynamic thresholding algorithm. The $IC_{50}$ point, as determined by the growth rate, is shown as a vertical green line. The threshold is a horizontal line between the control GRH and the largest magnitude change. The cell MIC is where the GRH curve first crosses this threshold. This generally corresponds well with $IC_{50}$ point as determined by the growth rate. Note that the signal from individual repeats is too noisy to perform this analysis accurately.

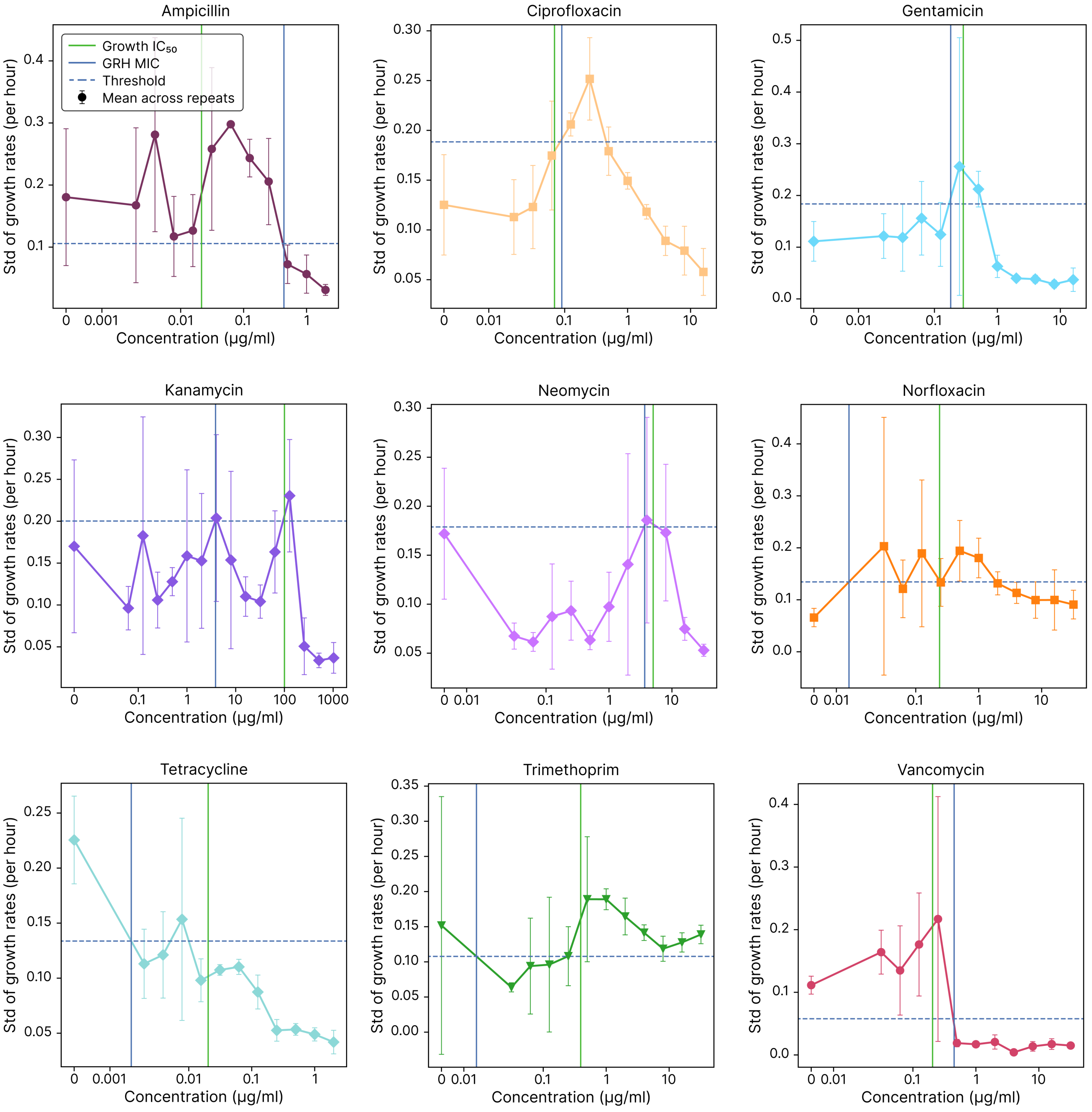


Figure S14: Showing how GRH can be used as a metric to determine MIC for *S. aureus*, see legend of fig. S13 for details.

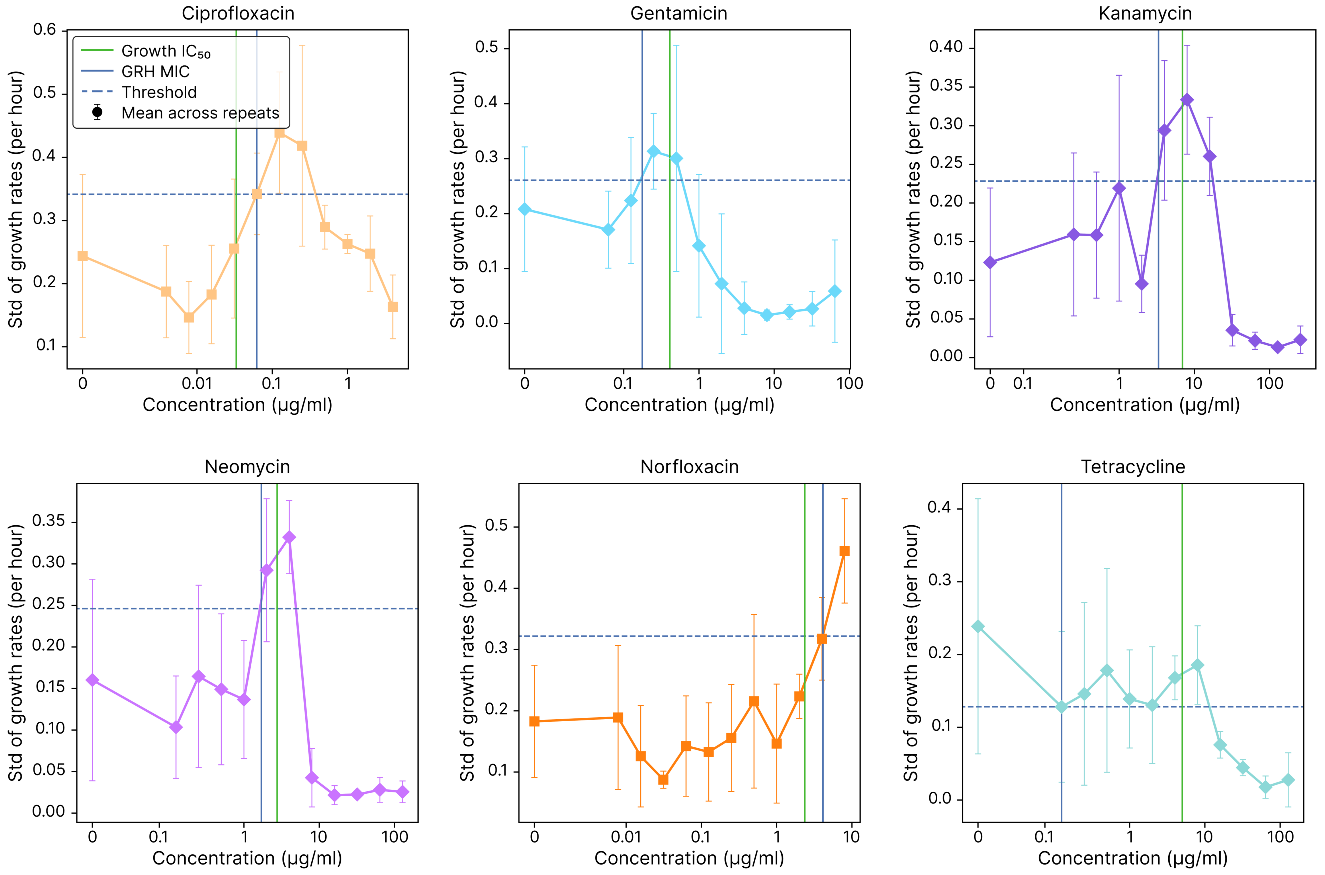


Figure S15: Showing how GRH can be used as a metric to determine MIC for *P. aeruginosa*, see legend of fig. S13 for details.

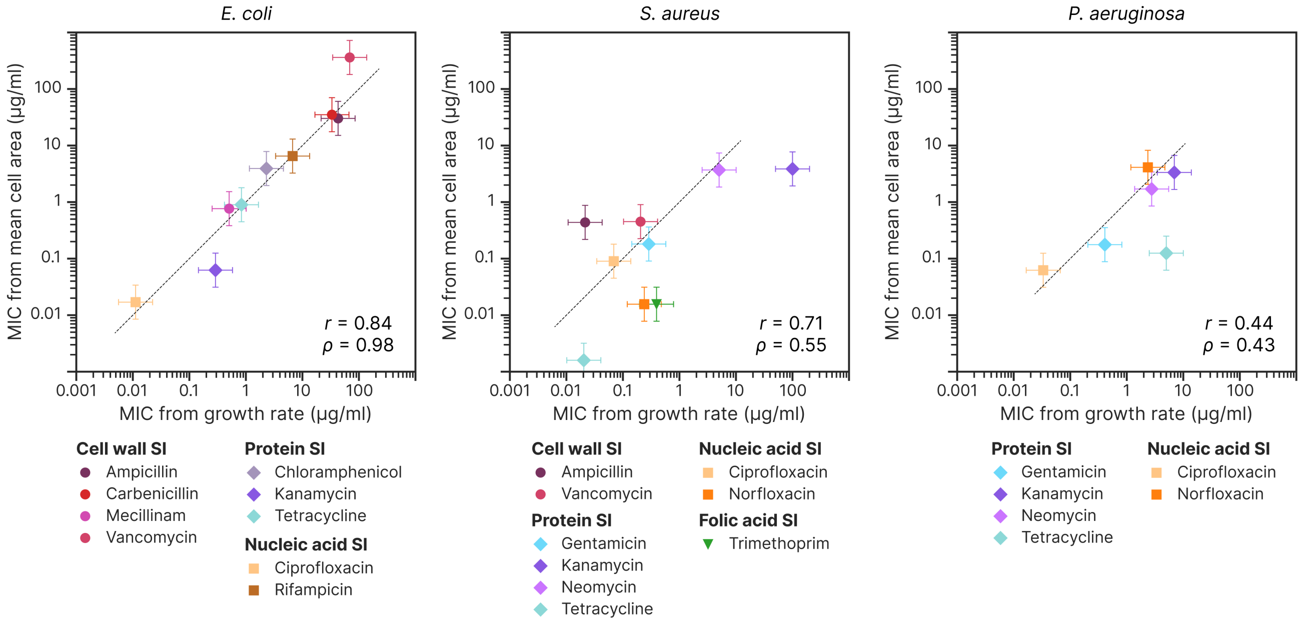


Figure S16: A summary of how well MIC from thresholding based on GRH corresponds to MIC based on growth rate (here represented by the $IC_{50}$ concentration, not $IC_{90}$). The x-axis shows MIC values based on growth rate, and the y-axis shows MIC values based on GRH. The dashed line represents the x=y. Pearson's correlation coefficient $r$ and Spearman's rank correlation coefficient $\rho$ are included for each species.

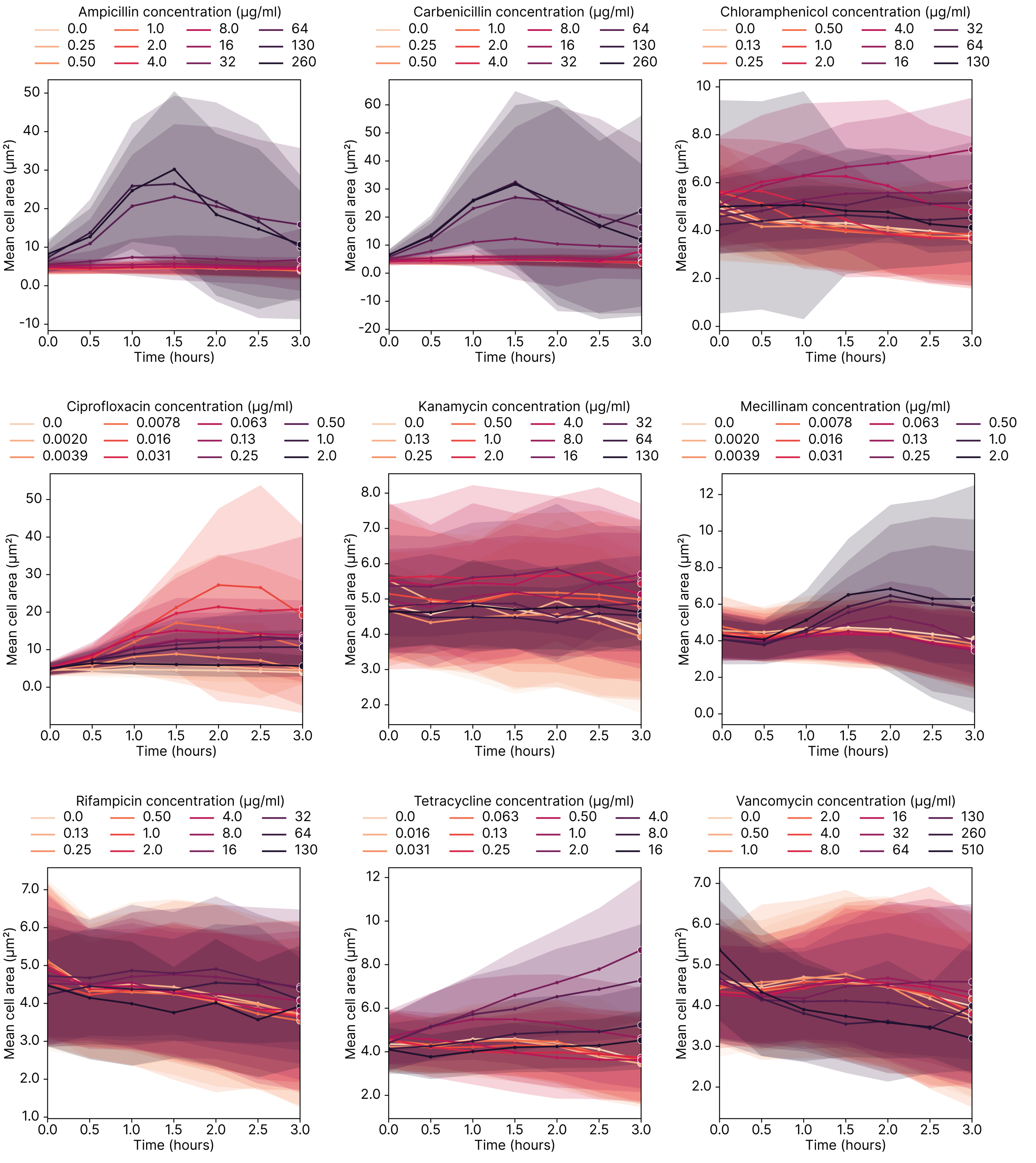


Figure S17: Showing how cell area changes over time when exposed to various concentrations of the antibiotics for *E. coli*. Initially, all bacteria have the same morphology as they are introduced to the pads with antibiotics. As time progresses, many antibiotics cause a significant change in morphology for some concentrations. Generally, the largest differences are apparent after 2 hours of growth. Each line represents the mean area per cell for a given antibiotic concentration, and the shaded area represents the standard deviation with data originating from four repeats. Darker colours correspond to higher antibiotic concentration, and data points are binned to the nearest 30 minutes.

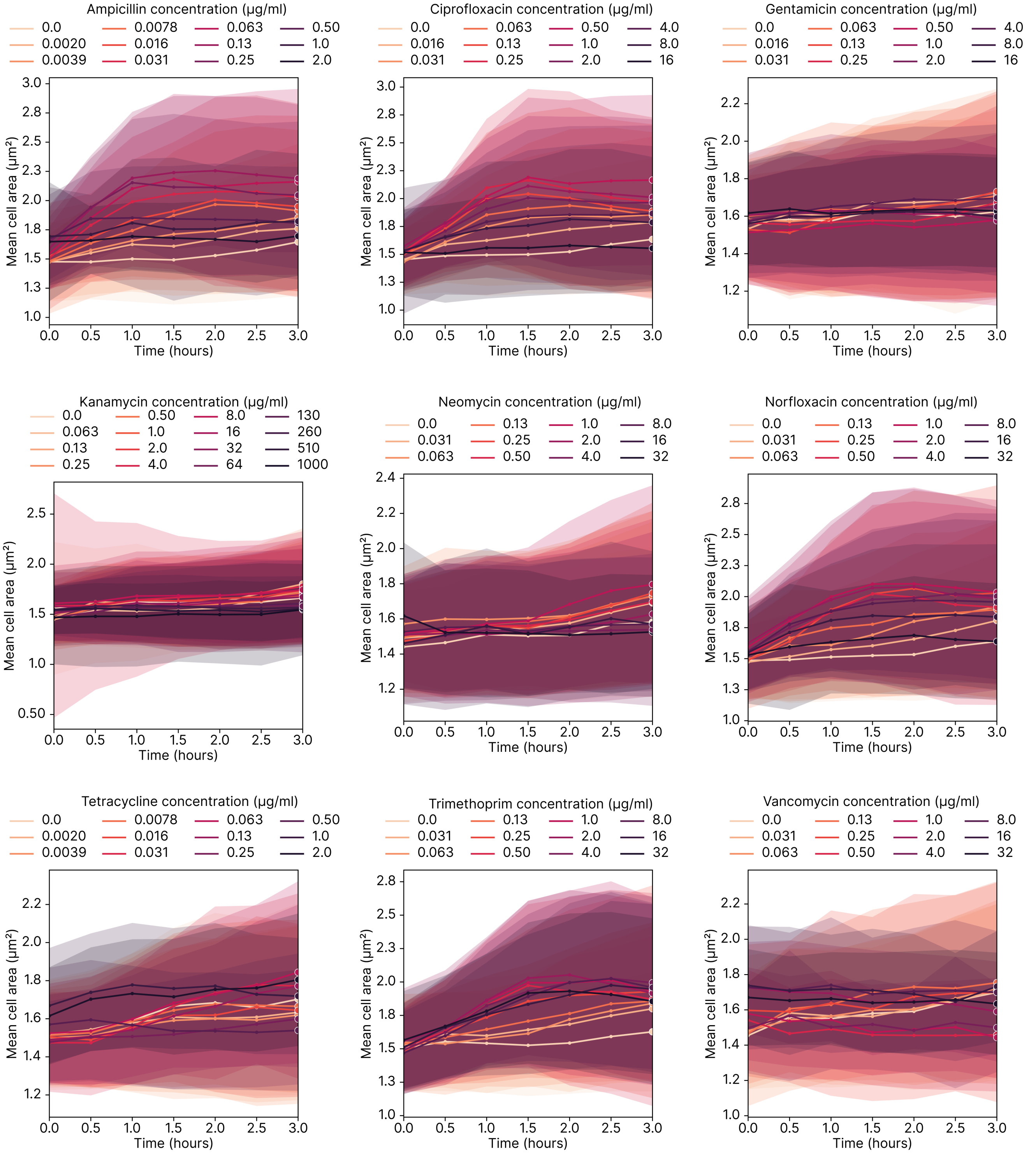


Figure S18: Showing how cell area changes over time when exposed to various concentrations of the antibiotics for *S. aureus*, see fig. S17 for details.

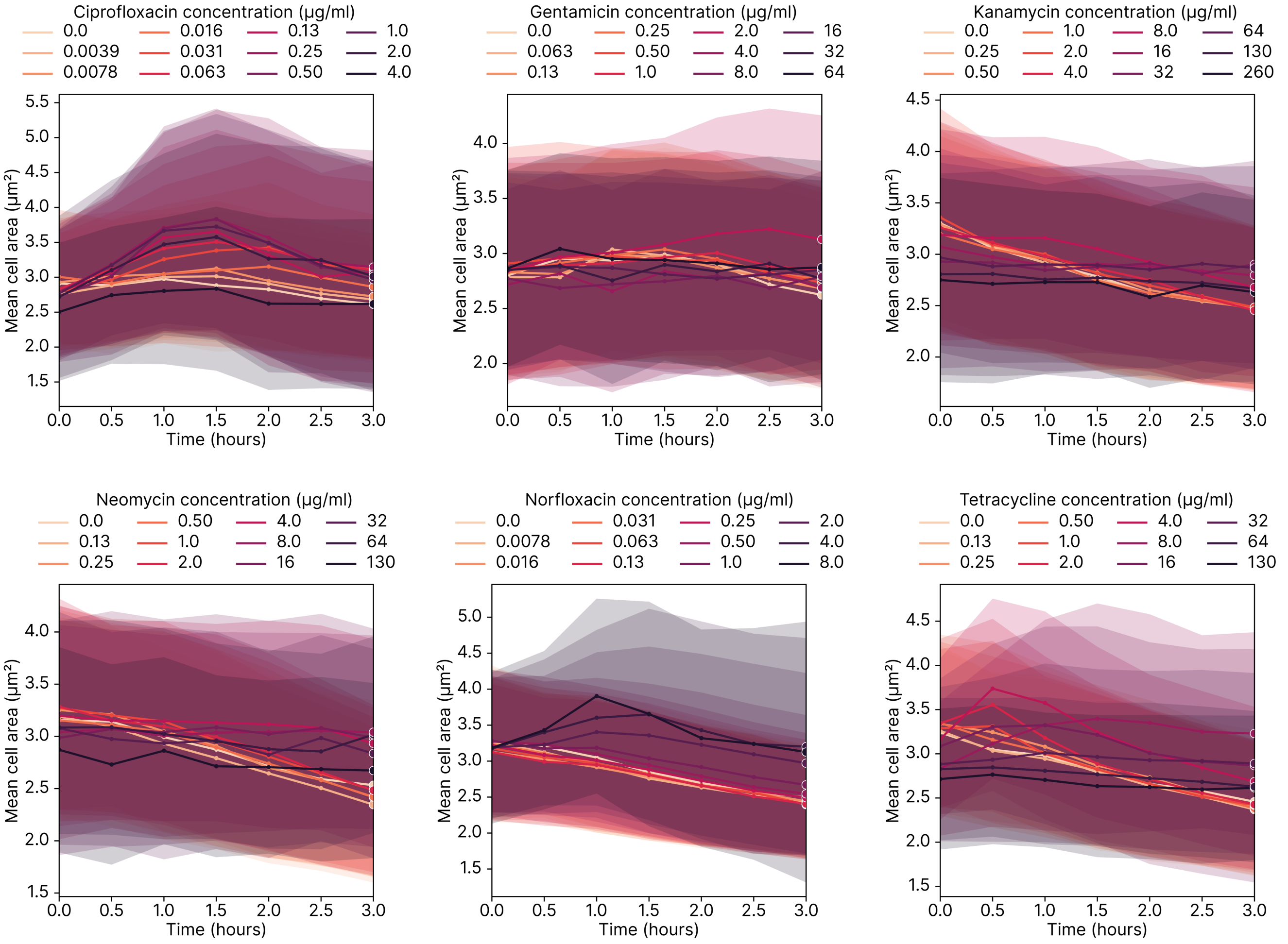


Figure S19: Showing how cell area changes over time when exposed to various concentrations of the antibiotics for *P. aeruginosa*, see fig. S17 for details.

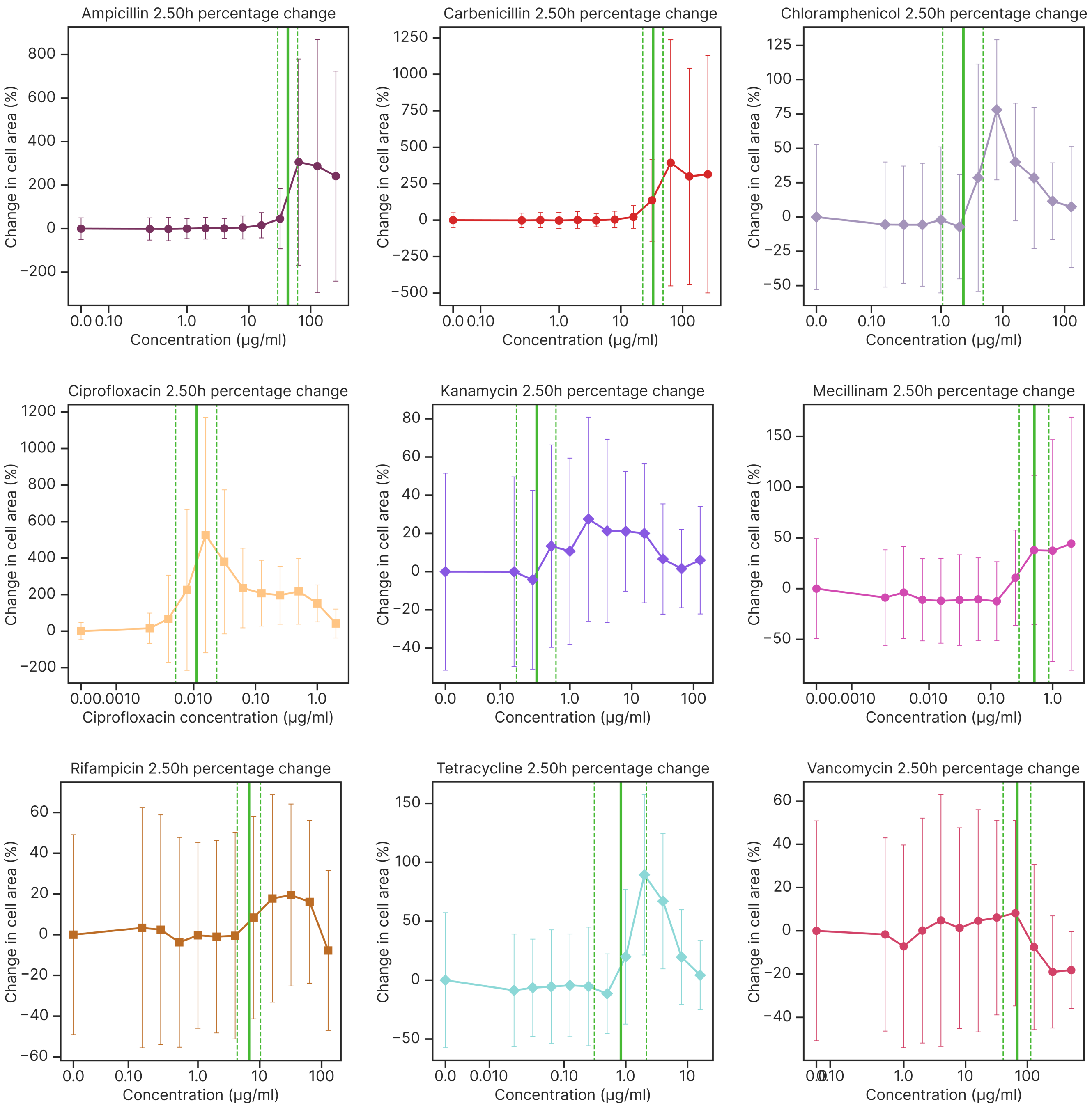


Figure S20: Showing how the mean cell areas change with antibiotic concentration for *E. coli*. The points represent the mean cell area from the repeats, and the error bars relate the standard deviation in cell areas based on data from the 2.5-hour mark. The vertical lines represent $IC_{10}$, $IC_{50}$, $IC_{90}$ respectively.

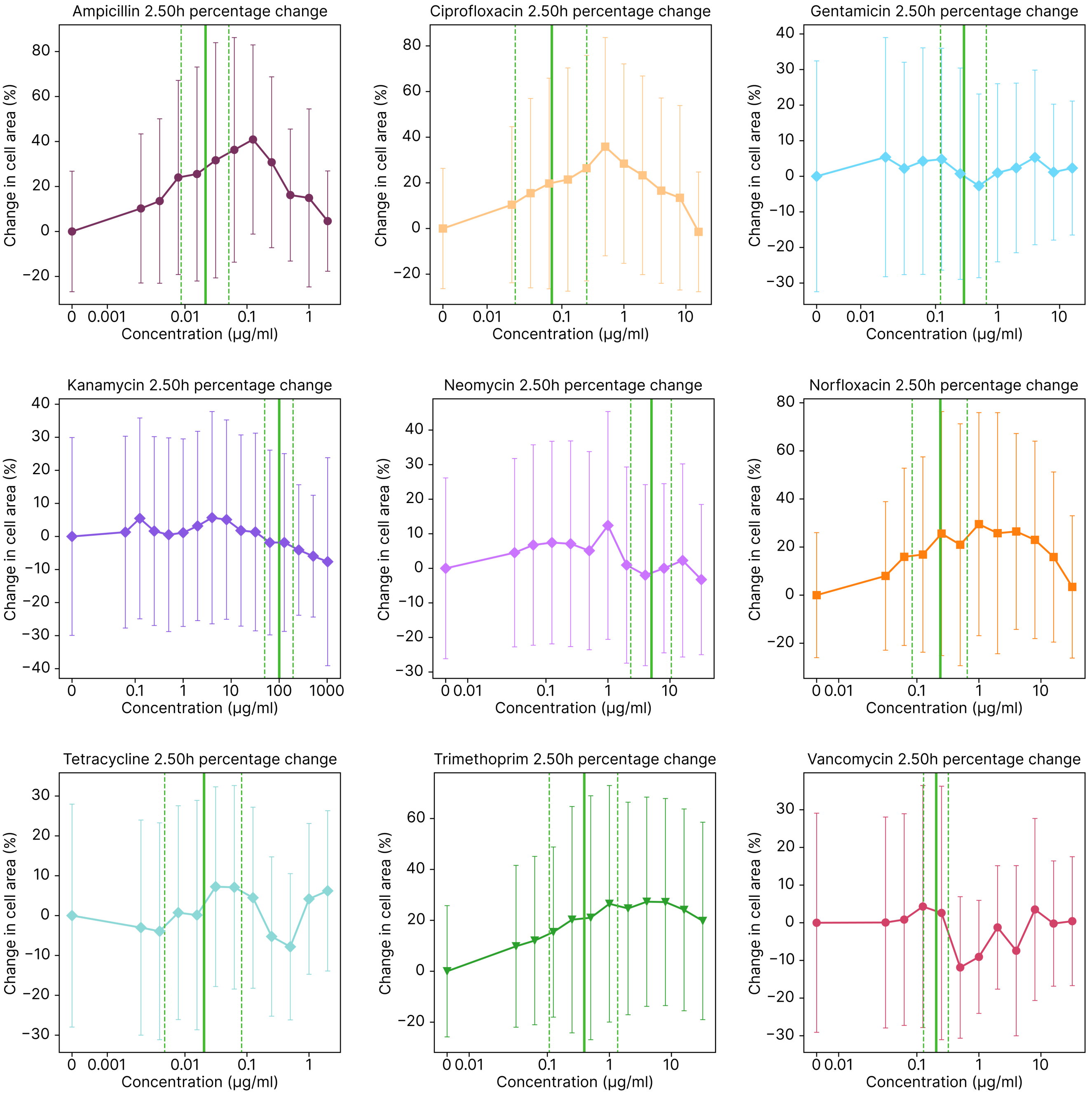


Figure S21: Showing how the mean cell areas change with antibiotic concentration for *S. aureus*, see fig. S20 for details.

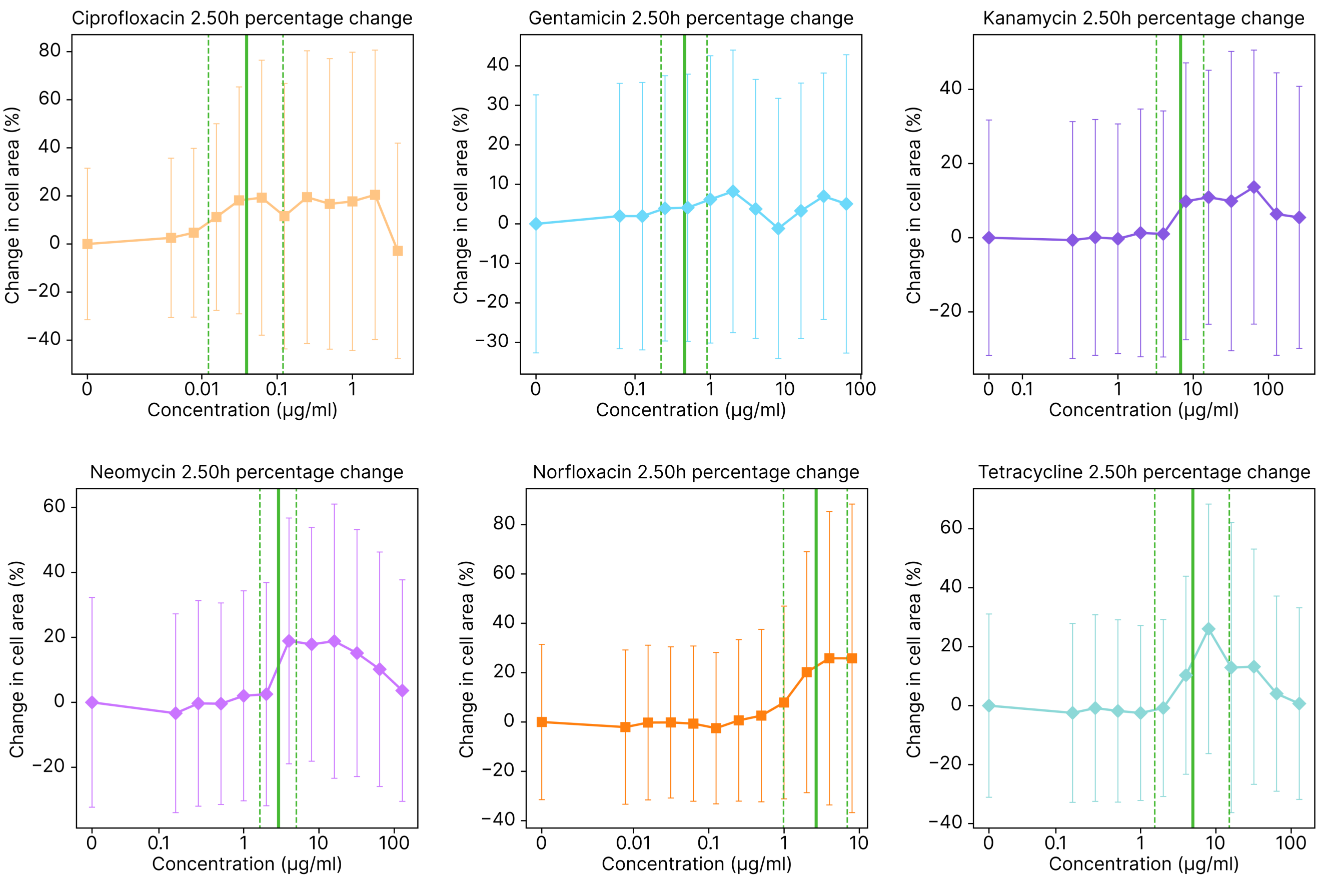


Figure S22: Showing how the mean cell areas change with antibiotic concentration for *P. aeruginosa*, see fig. S20 for details.

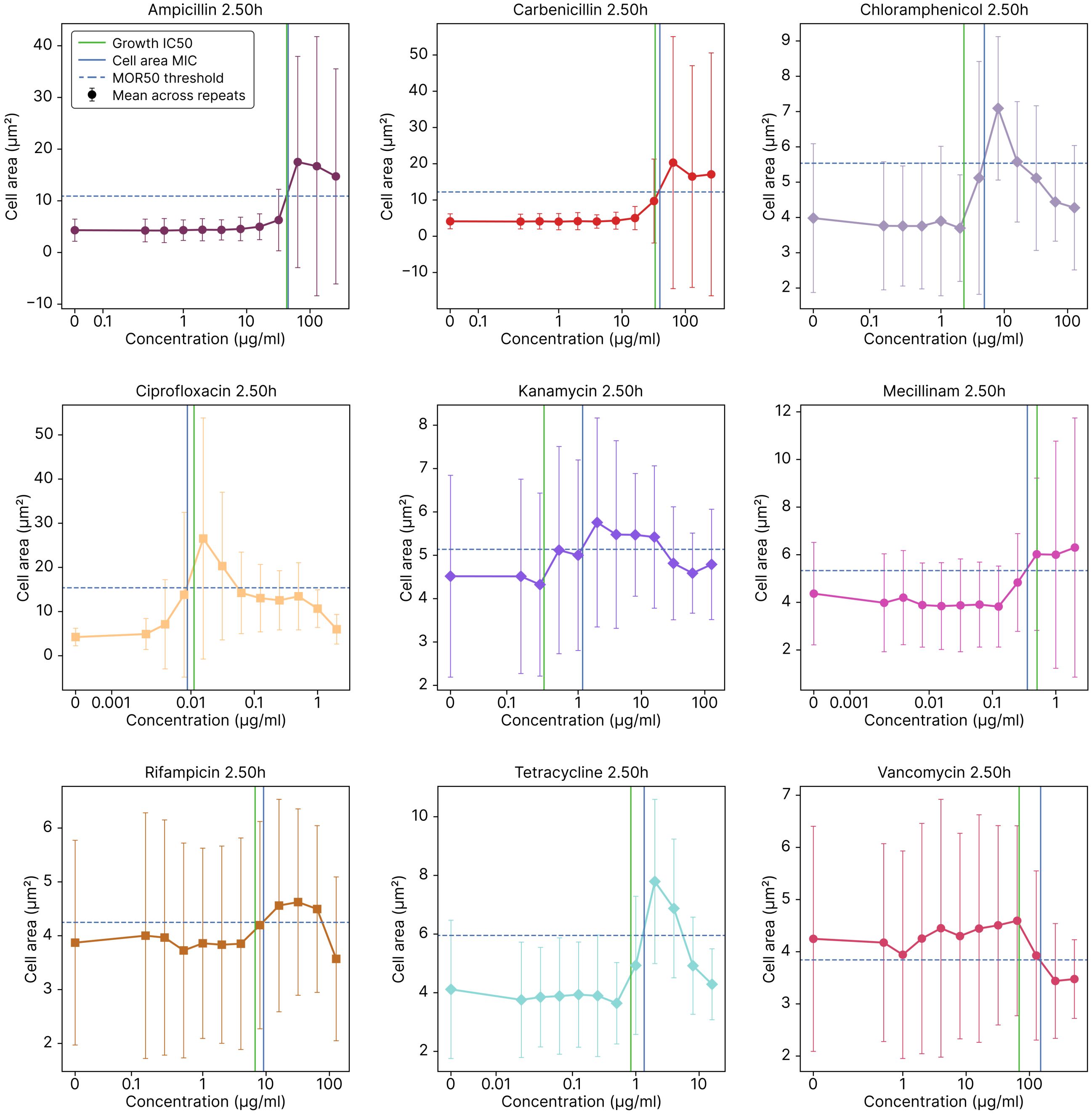


Figure S23: Showing how the mean cell areas change with antibiotic concentration with MOR50 information for *E. coli*. The points represent the mean cell area from the repeats, and the error bars relate the standard deviation in cell areas based on data from the 2.5-hour mark. The MOR50 threshold is halfway between the area with no antibiotic present and the maximum area change and is shown as a horizontal line. The cell MIC is where the area curve first crosses this threshold. This corresponds very closely with the $IC_{50}$ point as determined by the growth rate, represented by the vertical green line.

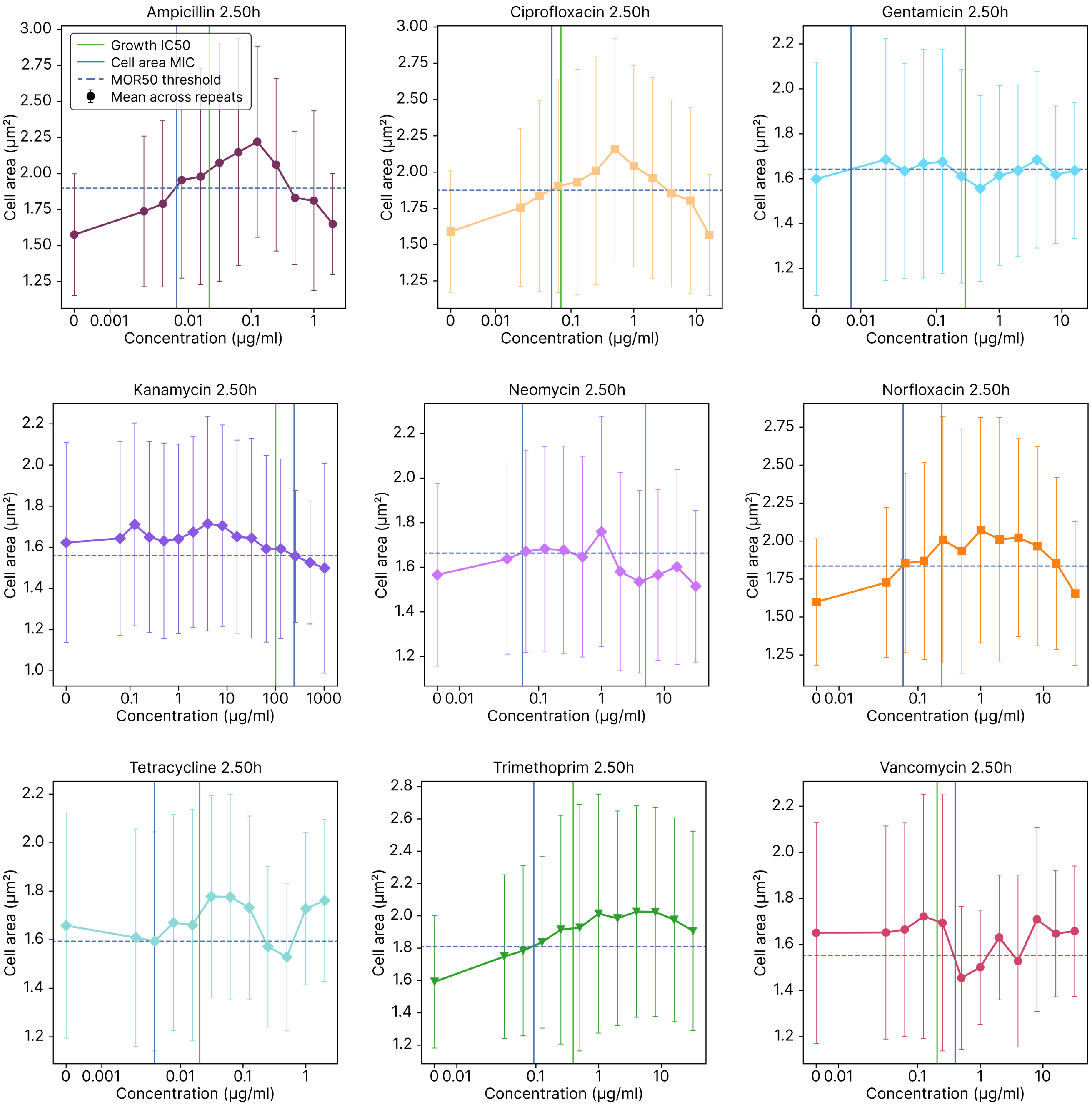


Figure S24: Showing how the mean cell areas change with antibiotic concentration with MOR50 information for *S. aureus*, see fig. S23 for details.

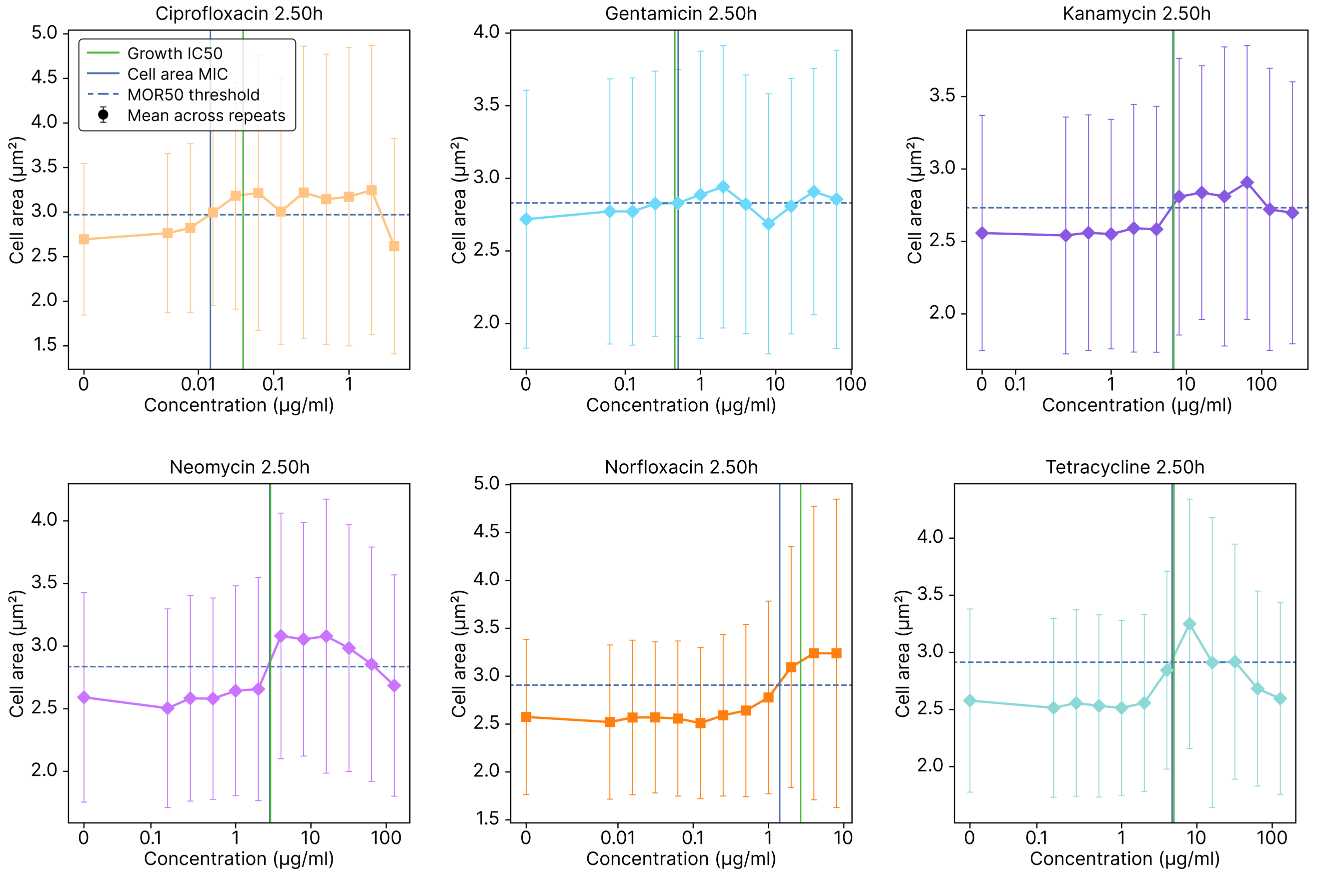


Figure S25: Showing how the mean cell areas change with antibiotic concentration with MOR50 information for *P. aeruginosa*, see fig. S23 for details.

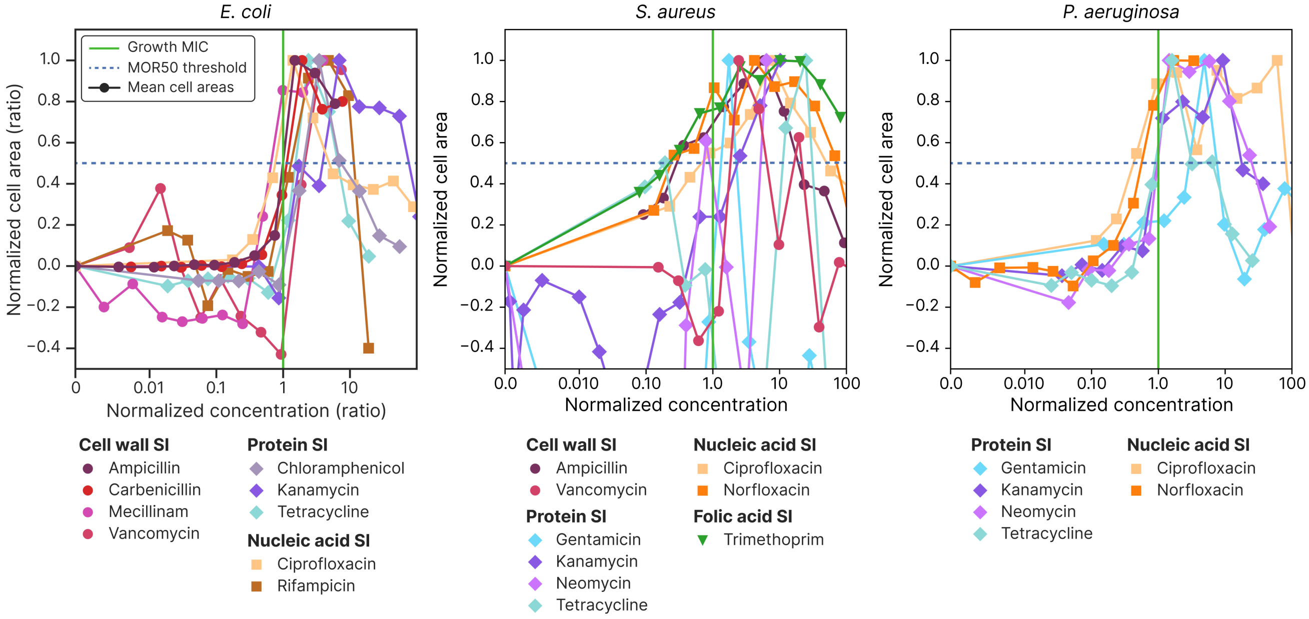


Figure S26: Showing how cell area changes depend on antibiotic concentration for *E. coli*, *S. aureus* and *P. aeruginosa*. The plot for *E. coli* is also shown in the main text but repeated here for completeness. For each species, curves are shown for all the tested antibiotics after 2.5 hours of incubation with the antibiotics. The antibiotic concentrations are normalised to growth MIC, and the cell area is normalised so that the no-antibiotic area is 0 and the maximum change in cell area is 1 (which for vancomycin entails inverting the area). The MOR50 threshold is shown as a horizontal line at 50% change, and the cell MIC is where the area curve first crosses this threshold. All the antibiotics produce a morphology change around the growth MIC. The degree of change varies strongly between antibiotics, which is here made apparent through the noise present in the area signal for antibiotics that induces a small change in morphology. The antibiotics are categorised by action mechanisms as inhibiting cell wall synthesis, protein synthesis, and nucleic acid synthesis, in addition to a negative control where no antibiotic was used.

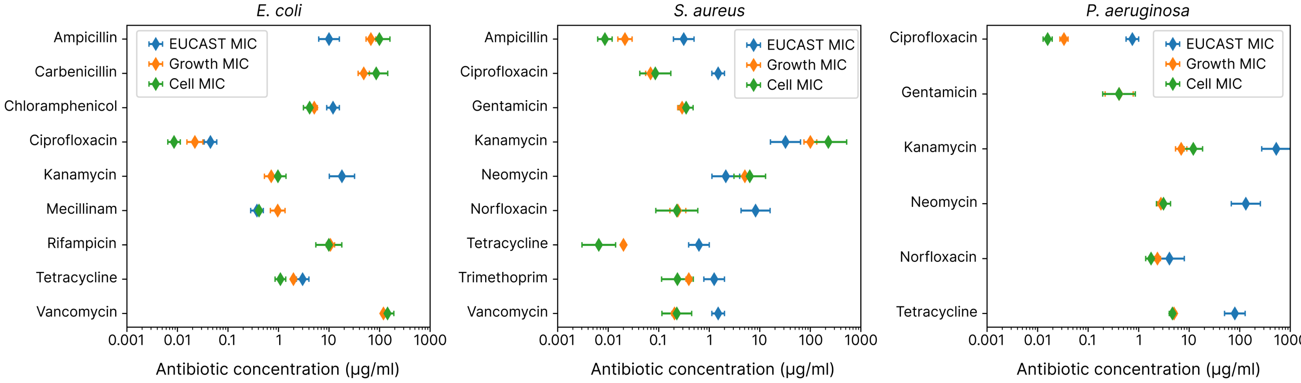


Figure S27: Showing how MIC measurements compare between EUCAST tabulated data from table S3, measurements based on growth rate measured on MAP platform, and MOR50 metric on MAP platform for *E. coli*, *S. aureus* and *P. aeruginosa*.
